# Mitochondrial-encoded complex I impairment induces a targetable dependency on aerobic fermentation in Hürthle cell carcinoma of the thyroid

**DOI:** 10.1101/2022.08.16.504162

**Authors:** Anderson R Frank, Vicky Li, Spencer D Shelton, Jiwoong Kim, Gordon M Stott, Leonard Neckers, Yang Xie, Noelle S Williams, Prashant Mishra, David G McFadden

**Affiliations:** Department of Internal Medicine, Division of Endocrinology, University of Texas Southwestern Medical Center, Dallas, TX 75390, USA; Department of Biochemistry, University of Texas Southwestern Medical Center, Dallas, TX 75390, USA; Children’s Medical Center Research Institute, University of Texas Southwestern Medical Center, Dallas, TX 75390, USA; Department of Population and Data Sciences, University of Texas Southwestern Medical Center, Dallas, TX 75390, USA; Leidos Biomedical Research Inc., Frederick National Laboratory for Cancer Research, Frederick, MD 24060, USA; Urologic Oncology Branch, Center for Cancer Research, National Cancer Institute, NIH, Bethesda, MD 20892, USA; Harold C. Simmons Comprehensive Cancer Center, University of Texas Southwestern Medical Center, Dallas, TX 75390, USA; Deparment of Pediatrics, University of Texas Southwestern Medical Center, Dallas, TX 75390, USA; Program in Molecular Medicine, University of Texas Southwestern Medical Center, Dallas, TX 75390, USA

## Abstract

A metabolic hallmark of cancer identified by Warburg is the increased consumption of glucose and secretion of lactate, even in the presence of oxygen. Although many tumors exhibit increased glycolytic activity, most forms of cancer rely on mitochondrial respiration for tumor growth. We report here that Hürthle cell carcinoma of the thyroid (HTC) models harboring mitochondrial DNA-encoded defects in complex I of the mitochondrial electron transport chain exhibit impaired respiration and alterations in glucose metabolism. CRISPR/Cas9 pooled screening identified glycolytic enzymes as selectively essential in complex I-mutant HTC cells. We demonstrate in cultured cells and a PDX model that small molecule inhibitors of lactate dehydrogenase selectively induce an ATP crisis and cell death in HTC. This work demonstrates that complex I loss exposes fermentation as a therapeutic target in HTC and has implications for other tumors bearing mutations that irreversibly damage mitochondrial respiration.

## INTRODUCTION

The mitochondrial electron transport chain (ETC) utilizes a series of multiprotein complexes to transfer electrons from reduced species to terminal electron acceptors and establishes a proton gradient across the mitochondrial inner membrane (Vafai and Mootha, 2012; Vercellino and Sazanov, 2022). ATP synthase harnesses the resulting proton gradient during oxidative phosphorylation to efficiently generate cellular ATP from various fuel sources. Beyond ATP production, mitochondrial ETC activity and respiration affect cellular and mitochondrial health by regulating cellular redox status (Titov et al., 2016), supporting aspartate biosynthesis (Birsoy et al., 2015; Sullivan et al., 2015), and sustaining mitochondrial protein import (Neupert and Herrmann, 2007). Under conditions of limiting oxygen, or impaired mitochondrial ETC function, cellular energy production shifts to glycolysis. To support glycolytic flux under these conditions, the product of glycolysis, pyruvate, is converted to lactate in a process known as fermentation and subsequently exported from the cell. The concerted actions of glycolysis and fermentation thus allow cells to meet the energy demands for survival and proliferation when oxygen is limiting, or oxidative phosphorylation is disrupted.

The observation by Warburg that tumors exhibit high rates of glucose consumption and lactate secretion in the presence of oxygen led him to hypothesize that these metabolic changes stem from respiratory deficiencies (Koppenol et al., 2011; Warburg, 1956). Indeed, a longstanding observation is that cancer cell lines and tumors exhibit increased glucose uptake and concentration relative to normal tissues, and these differences in glucose metabolism form the basis of ^18^FDG-PET imaging (Luengo et al., 2017). Consistent with Warburg’s early observations, cultured cancer cells perform glycolysis and fermentation even in the presence of oxygen (aerobic glycolysis, aerobic fermentation, or the Warburg effect), converting much of this glucose to lactate, and recent work has proposed that aerobic glycolysis allows proliferating cells to meet NAD^+^ demands (Luengo et al., 2021).

Multiple lines of evidence, including studies directly assessing tumor metabolism in patients, however, have demonstrated that most cancers are respiration-competent and readily catabolize fuels via the tricarboxylic acid (TCA) cycle (Faubert et al., 2017; Hensley et al., 2016; Johnston et al., 2021). Additionally, mitochondrial DNA (mtDNA) replication, a process required for production of a functional respiratory chain, is necessary for tumorigenesis and tumor cells subjected to mtDNA depletion go so far as to acquire healthy mitochondria *in vivo* to recover respiratory capacity (Tan et al., 2015; Weinberg et al., 2010). Multiple studies have also demonstrated that pharmacologic inhibition of mitochondrial ETC activity impairs tumor cell growth *in vitro* and *in vivo*, and next-generation ETC inhibitors are currently under investigation as anti-cancer therapeutics (Evans et al., 2021; Molina et al., 2018; Wheaton et al., 2014). Thus, while Warburg’s observation that cancers readily perform glycolysis and fermentation has been repeatedly verified, his hypothesis that cancers possess respiratory deficiencies and that these deficiencies are required for oncogenesis has largely been proven incorrect.

Mitochondrial ETC complexes are unique in that the individual protein subunits are encoded by genes in both the nuclear and mitochondrial genomes. Core subunits of complexes I (NADH dehydrogenase), III (cytochrome *bc*_*1*_ oxidoreductase), IV (cytochrome *c* oxidase), and V (ATP synthase) are mtDNA-encoded (Vafai and Mootha, 2012; Vercellino and Sazanov, 2022), and disruptive mtDNA mutations affecting these subunits have been linked to several mitochondrial diseases (Vafai and Mootha, 2012). Although somatic mtDNA mutations are common in cancer, analyses of large tumor cohorts from diverse cancer types have highlighted that most cancers select against loss-of-function mtDNA mutations (Ju et al., 2014; Tasdogan et al., 2020; Yuan et al., 2020). Selection against deleterious mtDNA mutations supports the notion that a functional respiratory chain is required to meet the metabolic demands of a proliferating tumor. Pan-cancer sequencing studies have highlighted exceptions including colon, kidney, and thyroid cancers that harbor recurrent high allelic fraction truncating mtDNA mutations (Yuan *et al*., 2020). In addition, sequencing efforts focused on oncocytic tumors (oncocytomas) have demonstrated that these tumors display significant enrichment of loss-of-function mtDNA mutations (Ganly et al., 2018; Gasparre et al., 2007; Gopal et al., 2018a; Gopal et al., 2018b). Oncocytomas arise in diverse sites including the salivary gland, kidney, and thyroid gland, and are defined by tumor cells exhibiting substantial mitochondrial accumulation. Electron microscopy studies of oncocytic thyroid tumors have revealed that these mitochondria are distended and exhibit irregular cristae patterns, suggestive of mitochondrial dysfunction (Valenta et al., 1974). The disruptive mtDNA mutations identified in these studies provide a possible explanation for the observed changes in mitochondrial abundance and morphology.

Hürthle cell carcinoma of the thyroid (HTC, also called oncocytic carcinoma of the thyroid) is an oncocytic form of thyroid cancer characterized by mitochondrial accumulation, increased glucose uptake, poor radioiodine concentration, and distinct patterns of metastatic spread (Ganly and McFadden, 2019; McFadden and Sadow, 2021). We and others (Ganly *et al*., 2018; Gopal *et al*., 2018b) have reported that mtDNA-encoded mutations in complex I of the mitochondrial ETC occur in approximately 60% of HTC tumors. Missense and loss-of-function complex I mtDNA mutations are significantly enriched in HTC and retained during disease progression, suggesting a driver role for these mutations. While these genetically encoded respiratory defects would suggest that HTC represents a true “Warburgian” tumor, the impact of these mutations on mitochondrial ETC function and cellular metabolism in the setting of HTC remains incompletely characterized. Recent work has highlighted how an understanding of genetically driven alterations in metabolism can be used to identify synthetic vulnerabilities in several cancers (Dang et al., 2009; Mardis et al., 2009; Rohle et al., 2013; Romero et al., 2017; Yan et al., 2009). We reasoned that investigation of the functional consequences of mtDNA mutations in HTC would expose new therapeutic vulnerabilities. In this study, we demonstrate that HTC cells with disruptive complex I mtDNA mutations exhibit mitochondrial ETC dysfunction and defects in central carbon metabolism. Using CRISPR/Cas9 screening and pharmacologic tools, we demonstrate that complex I-mutant HTC cells are strictly reliant on glycolysis, and that fermentation represents a therapeutically tractable liability in complex I-mutant HTC.

## RESULTS

### Patient-derived HTC models exhibit complex I impairment

We identified two patient-derived HTC models to facilitate study of mtDNA-encoded complex I mutations in HTC. The XTC.UC1 cell line, which is the only reported HTC model, was established from an HTC metastasis to breast, and prior studies reported deleterious mtDNA mutations in the complex I gene *MT-ND1*, in addition to chromosome level loss-of-heterozygosity (LOH) across multiple chromosomes (Bonora et al., 2006; Corver et al., 2018; Zielke et al., 1998). We acquired XTC.UC1 and confirmed the *MT-ND1* insertion (ChrM:3571C>CC; p.Leu89_FS_*12) in XTC.UC1 using Sanger sequencing; however, we observed both wild-type and mutant *MT-ND1* traces, suggesting a heteroplasmic shift in the XTC.UC1 mtDNA genotype (Figure S1N) (Bonora *et al*., 2006). We also identified a patient-derived xenograft (PDX) derived from a poorly-differentiated HTC at the National Cancer Institute Patient-Derived Models Repository (www.pdmr.cancer.gov/), NCI-248138-237-R, hereafter NCI-237-R^PDX^. We downloaded exome sequencing data from NCI-237-R^PDX^ available from the PDMR and analyzed on- and off-target sequencing reads to establish a genomic profile for NCI-237-R^PDX^. Our LOH analysis pipeline identified widespread LOH across every chromosome except Chr 7 and Chr 20, consistent with the extreme chromosome losses we and others previously reported in large cohorts of HTC (Figure 1A) (Ganly et al., 2018; Gopal et al., 2018).

**Figure 1:**
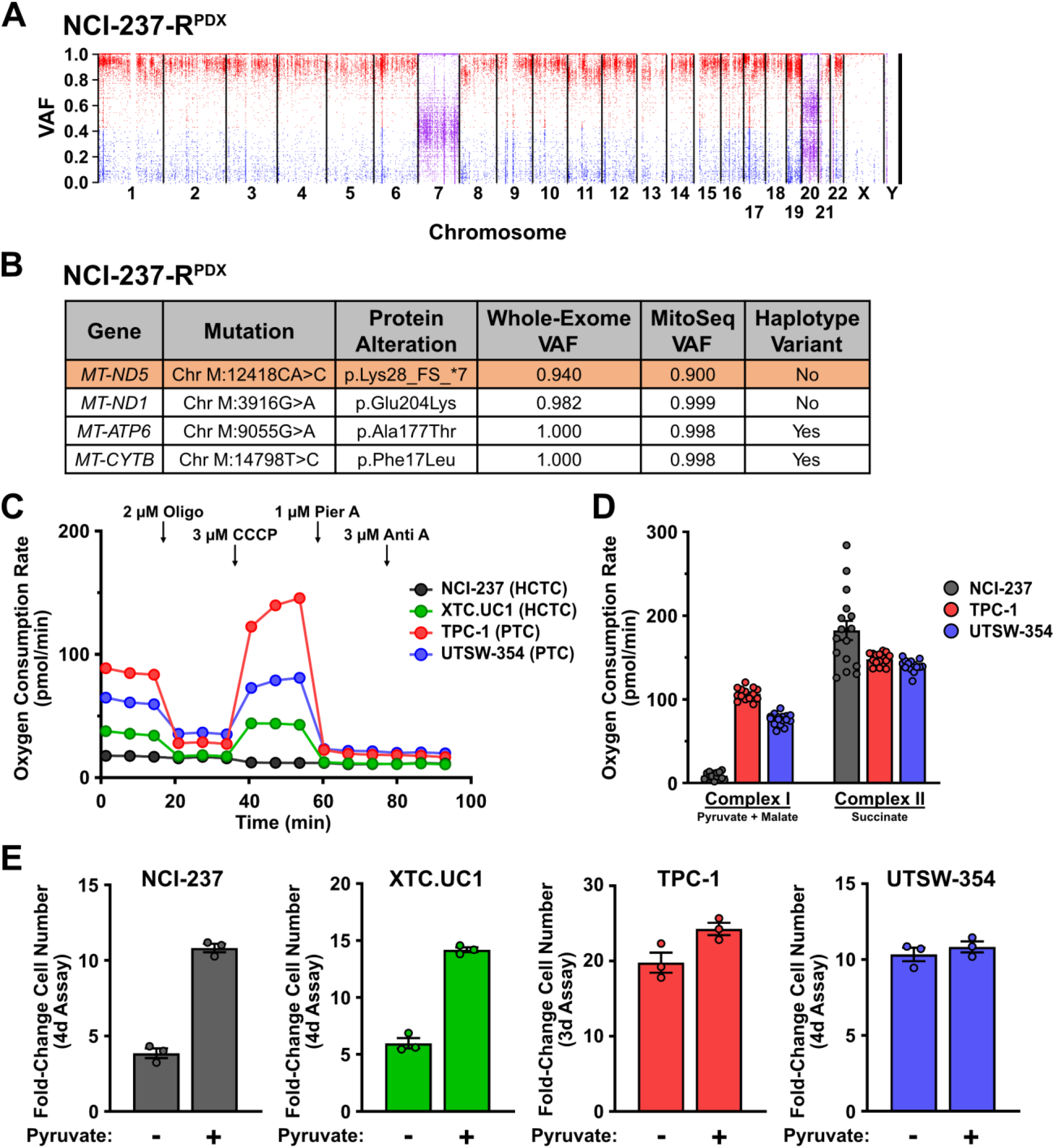
Patient-derived HTC models exhibit complex I impairment. A) Variant allele fraction (VAF) plot for NCI-237-R^PDX^. Variants with VAF > 0.5 are in red; variants with VAF < 0.5 are in blue. B) Table of NCI-237-R^PDX^ nonsynonymous mtDNA mutations identified through whole-exome sequencing and targeted mtDNA sequencing (MitoSeq). C) Oxygen consumption rates for intact cells; *n* = 6-8 replicates. Oligo = oligomycin; CCCP = carbonyl cyanide 3-chlorophenylhydrazone; Pier A = piericidin A; Anti A = antimycin A. D) Oxygen consumption rates for permeabilized cells; *n* = 16 replicates and points represent individual wells. E) Fold-change in cell number of indicated cell lines in media -/+ 100 μM pyruvate. *n* = 3 replicates. Data are plotted as mean ± SEM of indicated number of replicates.

Review of off-target exome sequencing reads that mapped to mtDNA demonstrated a coverage depth of 212X, which was sufficient to identify mutations. To further improve resolution and fidelity of mtDNA mutation profiling, we performed targeted mtDNA sequencing from NCI-237-R^PDX^ samples collected after establishing the model in our laboratory (Figure S1A). Targeted mtDNA sequencing yielded a mean coverage depth of 7,239X (Wang et al., 2022a). Our analysis pipelines identified two non-haplotype mtDNA mutations of interest: a near-homoplasmic deleterious mutation in *MT-ND5* resulting in early termination (ChrM:12418CA>C; p.Lys28_FS_*7; VAF^WES^ = 0.940; VAF^MitoSeq^ = 0.900), as well as a near-homoplasmic missense mutation of unknown functional significance in *MT-ND1* resulting in a Glu to Lys substitution (ChrM:3916G>A; p.Glu204Lys; VAF^WES^ = 0.982; VAF^MitoSeq^ = 0.999) (Figure 1B and Table S1). We validated both mutations using Sanger sequencing (Figures S1B and S1C). These data demonstrate that NCI-237-R^PDX^ harbors a disruptive mtDNA mutation affecting complex I of the mitochondrial ETC.

Consistent with mutations reported by the PDMR, we identified nonsense mutations in the tumor suppressors *NF1* (Chr17:31261733C>T; p.Arg1534*) and *NF2* (Chr22:29671910C>T; p.Gln362*), as well as a series of deletions in exon 9 of the DNA methyltransferase, *DNMT3A* (Chr2:25247091 TTGTTGTACGTGGCCTGGT>TC; p.His355_FS_*32), that caused frameshifting and early termination. These mutations were validated by Sanger sequencing and, in line with the reported high allelic fraction and widespread LOH, appear homozygous (Figures S1D-S1F and Table S2). Based on these data, we conclude that NCI-237-R^PDX^ harbors the hallmark genetic features of HTC.

To enable detailed characterization of mitochondrial ETC function, we derived a cell line from a dissociated NCI-237-R^PDX^ tumor (the cell line is hereafter referred to as NCI-237^UTSW^). We determined that NCI-237^UTSW^ forms subcutaneous xenografts following re-implantation into immunodeficient mice and verified that key mitochondrial and nuclear genome mutations were preserved during cell line establishment (Figures S1G-S1L and Table S1).

To evaluate alterations in mitochondrial ETC function and metabolism in complex I-mutant HTC cell lines, we utilized a panel of thyroid cancer cell lines including NCI-237^UTSW^, XTC.UC1, TPC-1 (papillary thyroid carcinoma (PTC) cell line with RET/CCDC6 translocation), and UTSW-354 (PTC cell line with BRAF^V600E^ mutation) (Figure S1M). Because no descriptions of mtDNA mutations were available for TPC-1 and UTSW-354, we performed targeted mtDNA sequencing to identify potential deleterious mutations (mean coverage depth = 7,476X and 5,515X, respectively). Targeted mtDNA sequencing identified a low allelic fraction 1 base pair insertion in *MT-ND5* in TPC-1 (ChrM:12417A>AA; p.Asn27_FS_*31; VAF^MitoSeq^ = 0.02) and a heteroplasmic in-frame insertion in *MT-ND4* in UTSW-354 (ChrM:11866A>ACCC; p.Pro370Dup; VAF^MitoSeq^ = 0.65) (Table S1).

We assessed respiratory function in the panel of cell lines using Seahorse extracellular flux assays (Salabei et al., 2014). We observed that NCI-237^UTSW^ displayed dramatically reduced oxygen consumption that was not responsive to pharmacologic modulators of the ETC (Figure 1C). XTC.UC1 displayed approximately 40% oxygen consumption relative to TPC-1 and retained limited ETC-linked respiratory activity (Figure 1C). The *MT-ND1* mutation we validated in XTC.UC1 cells appeared heteroplasmic, suggesting that the fraction of wild-type mtDNA could encode limited respiratory capacity in these cells (Figure S1N). UTSW-354 cells exhibited slightly decreased oxygen consumption compared to TPC-1, and it is possible the in-frame MT-ND4^P370^ insertion we identified in UTSW-354 impacted complex I function. However, these cells did not display other features of impaired respiration (see below).

We next directly evaluated complex I function using permeabilized flux assays (Salabei *et al*., 2014). Incubation with pyruvate and malate (which stimulate complex I activity through NADH production) yielded minimal respiratory activity in NCI-237^UTSW^ cells (Figure 1D). In contrast, succinate (the substrate for complex II/succinate dehydrogenase) stimulated nearly equivalent respiratory activity across all cell lines, demonstrating intact complex II-IV function (Figure 1D). Combined, these data demonstrate that complex I-mutant HTC cell lines harboring mtDNA mutations at high allelic fraction exhibit impaired respiratory activity, most likely through a specific defect in complex I function.

Disruption of mitochondrial ETC activity induces widespread adaptations in central carbon metabolism, including alterations in glucose utilization (Mullen et al., 2011). To assess whether complex I-mutant HTC cell lines exhibit impaired glucose metabolism, we cultured cells with ^13^C-labeled glucose ([U-^13^C_6_] glucose) and tracked carbon fate using gas chromatography-mass spectrometry (GC-MS) (Figure S1O). All cell lines displayed similar enrichment of glucose-derived carbon in glycolysis-related metabolites. NCI-237^UTSW^, however, displayed minimal incorporation of glucose-derived carbon into the citrate pool via pyruvate dehydrogenase (PDH) and citrate synthase (CS) (Figure S1P). XTC.UC1 cells displayed similar citrate (m+2) levels to TPC-1 and UTSW-354, but analysis of downstream TCA cycle intermediates revealed low levels of (m+2)- and (m+3)-labeled species in XTC.UC1 cells, indicating reduced glucose-derived carbon movement through the TCA cycle in XTC.UC1 (Figure S1P).

Mitochondrial ETC dysfunction impairs NAD^+^ regeneration which leads to a requirement for exogenous pyruvate for cell growth. (Birsoy et al., 2015; King and Attardi, 1989; Sullivan et al., 2015; Titov et al., 2016). NCI-237^UTSW^ and XTC.UC1 displayed a proliferation defect when cultured in media lacking pyruvate (Figure 1E) whereas TPC-1 and UTSW-354 exhibited similar proliferation rates in pyruvate-free and pyruvate-replete media (Figure 1E). Based on these data, we conclude that patient-derived HTC models exhibit impaired mitochondrial ETC function and alterations in glucose metabolism consistent with damaging mtDNA mutations in complex I subunits.

### CRISPR/Cas9 screening identifies glycolysis as a metabolic dependency in complex I-mutant HTC

We hypothesized that complex I impairment in HTC cells required naturally evolved metabolic adaptations to support cellular proliferation, and that such adaptations might be potential therapeutic targets. To identify genes selectively essential for proliferation of complex I-deficient HTC cells, we utilized pooled CRISPR/Cas9 screening. We first performed a genome-wide CRISPR/Cas9 knockout screen using the Brunello single-guide RNA (sgRNA) library in NCI-237^UTSW^ and TPC-1 cells stably expressing Cas9. Cells were cultured for approximately 12 population doublings and sgRNA abundance in the initial and final cell populations was determined by next generation sequencing (Figures 2A and S2A). Rank ordering of genes based on the fold-change in sgRNA abundance (median of the 4 sgRNAs targeting a specific gene) identified multiple cell line-specific modifiers of cell growth (Figures S2B and S2C and Table S2). sgRNAs targeting positive (*ARNT* and *HIF1A)* and negative regulators (*VHL* and *EGLN1*) of the hypoxic stress response were enriched and depleted, respectively, in the NCI-237^UTSW^ screen (Figure S2B). Similarly, sgRNAs targeting *NFE2L2* (*NRF2*), the transcription factor controlling the oxidative stress response, were depleted in TPC-1 whereas sgRNAs targeting *KEAP1* and *CUL3*, which negatively regulate *NRF2* levels and activity, were enriched (Figure S2C). Together, these results suggested our genome-wide screens detected biologically linked essential genes.

**Figure 2:**
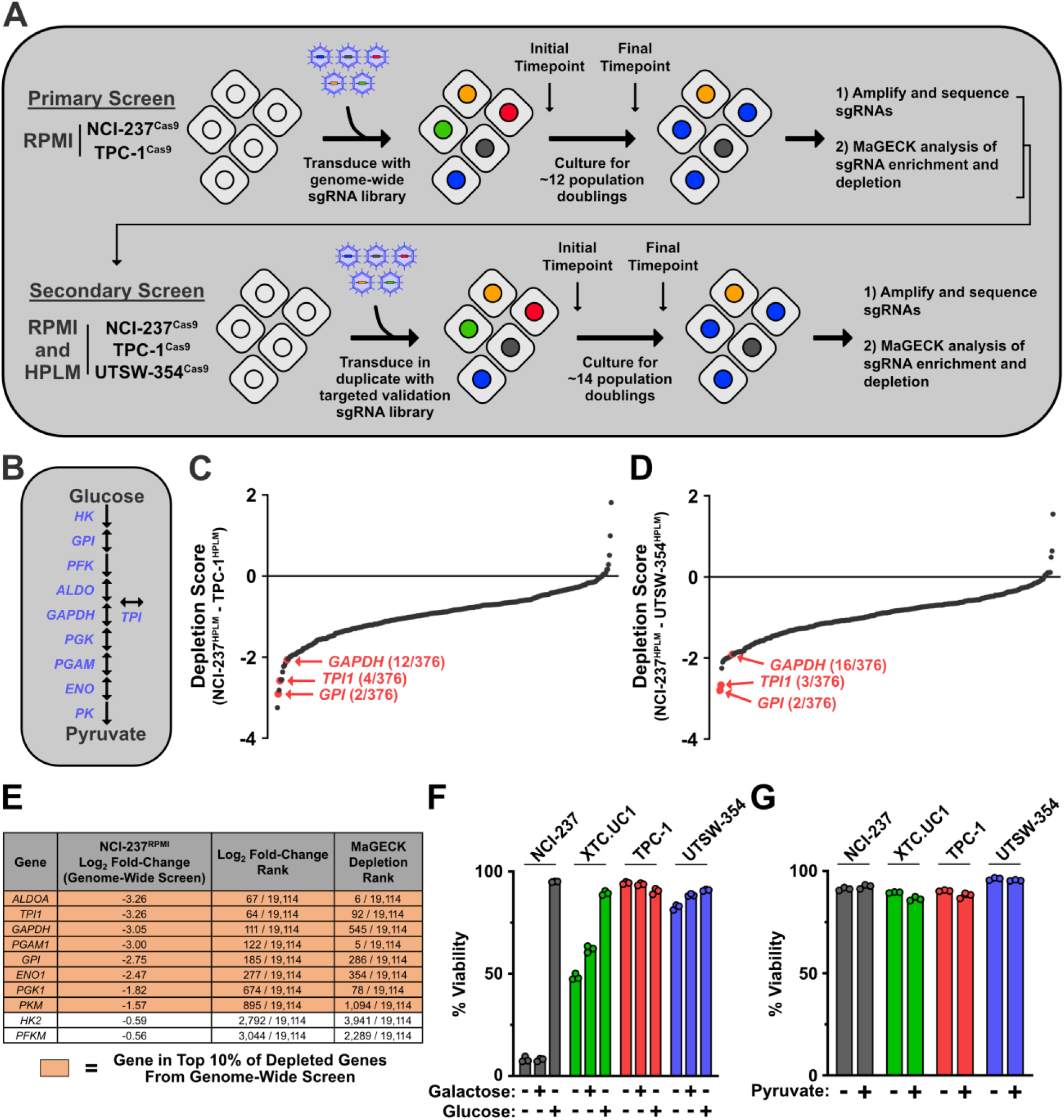
CRISPR/Cas9 screening identifies glycolysis as a metabolic dependency in complex I-mutant HTC. A) Schematic depicting CRISPR/Cas9 knockout screening pipeline. B) Schematic depicting the genes involved in glycolysis. C) Plot of genes from validation screen based on Depletion Score comparing NCI-237^HPLM^ and TPC-1^HPLM^. D) Plot of genes from validation screen based on Depletion Score comparing NCI-237^HPLM^ and UTSW-354^HPLM^. Depletion Scores for C) and D) are calculated as [(NCI-237^HPLM^ gene log_2_ fold-change) – (TPC-1^HPLM^ or UTSW-354^HPLM^ gene log_2_ fold-change)]. Gene log_2_ fold-change values reflect median log_2_ fold-change in sgRNA abundance and are based on 2 replicates for each cell line. E) Table of genes involved in glycolysis from NCI-237^RPMI^ genome-wide screen. Genes are listed according to log_2_ fold-change values. F) Viability of indicated cell lines cultured in media lacking glucose or galactose, or media containing 5 mM galactose or 5 mM glucose for 48 hours. G) Viability of indicated cell lines cultured in media -/+ 100 μM pyruvate for 48 hours. Data are plotted as mean ± SEM of 3 replicates.

We performed a targeted validation screen in which genes were selected based on three criteria: i) at least two-fold dropout in the primary NCI-237^UTSW^ genome-wide screen; ii) dropout was at least 1.5-fold greater in NCI-237^UTSW^ than TPC-1, and iii) the gene was not universally essential in The Cancer Dependency Map Project (DepMap) database (www.depmap.org/portal/). Three hundred fifty-three (353) genes met these criteria and the resulting library contained 1,579 sgRNAs targeting 376 genes (includes 23 control genes; 4 sgRNAs/gene and 75 non-targeting control sgRNAs taken from the Brunello library) (Table S2). To categorize genes whose effects were influenced by the cell culture media nutrient composition (Rossiter et al., 2021), we utilized NCI-237^UTSW^, TPC-1, and UTSW-354 cell lines cultured in both RPMI and physiologic human plasma like media (HPLM) (Cantor et al., 2017).

Cell lines stably expressing Cas9 were infected with the validation sgRNA library and cultured for approximately 14 population doublings and changes in sgRNA abundance were assessed by next generation sequencing of the sgRNA cassette from initial and final cell populations (Figures 2A, S2D, and S2E). Analysis of the validation screen revealed a positive correlation (R^2^ = 0.5398) between the fold-change in sgRNA abundance for NCI-237^RPMI^ and NCI-237^HPLM^ (Figure S2F). In addition, 303/353 and 229/353 genes displayed at least 2-fold dropout in NCI-237^UTSW^ cells cultured in RPMI and HPLM, respectively (Table S2). Together, these results suggested that the majority of genes validated in our targeted re-screen and that loss of most genes had similar effects on proliferation in RPMI and HPLM.

We compared gene dropout across cell lines and media conditions to identify genes whose loss selectively impaired NCI-237^UTSW^ proliferation. Consistent with our re-screen criteria, most genes showed selective depletion in NCI-237^UTSW^ (Figures 2C, 2D, S2G, and S2H). The glycolytic enzymes glucose-6-phosphate isomerase (*GPI)* and triose-phosphate isomerase 1 (*TPI1*) were both among the top 5 NCI-237^UTSW^-selective hits in the HPLM validation screen (Figures 2B-2D). In addition, glyceraldehyde-3-phosphate dehydrogenase (*GAPDH)*, which was included in the validation screen as a control for depletion across all cell lines, was in the top 20 NCI-237^UTSW^-selective hits (Figures 2B-2D). Glycolytic genes were also NCI-237^UTSW^-selective hits in the RPMI validation screen, but depletion was diminished compared to the HPLM validation screen (Figures S2G and S2H). This could be due to differential reliance on glycolysis for cells grown in traditional synthetic media versus physiologic media. Re-examination of the primary genome-wide screen revealed that genes encoding 8 of the 10 enzymes in glycolysis were in the top 10% of depleted genes in NCI-237^UTSW^ (Figure 2E). Interestingly, hexokinase and phosphofructokinase, which perform key irreversible steps in glycolysis, were not depleted (Figure 2E). These enzymes, however, are encoded by functionally redundant paralogs, which could explain why loss of a single gene did not significantly impair cell growth.

Glycolysis is a fundamental component of central carbon metabolism in all cells, and as such, several genes encoding glycolytic enzymes are “pan-essential” genes. Despite the universal essentiality of glycolysis, certain glycolytic genes, including *GPI* and *TPI1*, are not pan-essential, suggesting variable reliance on these enzymes. Why did these genes drop out in HTC cells compared to thyroid cancer cells with intact ETC function? We first tested if HTC cells responded differently to glucose withdrawal or galactose-containing media (Arroyo et al., 2016; Ditta et al., 1976; Robinson et al., 1992). We observed impaired proliferation of all cell lines in glucose-free or galactose-containing media irrespective of respiratory status (Figures S2I). However, NCI-237^UTSW^ and XTC.UC1 displayed reduced viability when cultured in glucose-free or galactose-containing media whereas TPC-1 and UTSW-354 remained viable (Figure 2F). XTC.UC1 viability loss was intermediate of NCI-237^UTSW^ and respiration-competent cells (Figure 2F), which is possibly a result of residual respiratory capacity (Figure 1A). Culture in pyruvate-free media, which selectively impaired complex I-mutant HTC cell proliferation, had no effect on cell viability, suggesting the metabolites linked to glucose and pyruvate availability have differential roles in cell proliferation and viability (Figure 2G). The depletion of genes encoding glycolytic enzymes in CRISPR screens and the observation of cell death following glucose withdrawal raised the possibility that a therapeutic window might exist for targeting glycolysis in HTC in the highly sensitized context of irreversible genetically encoded complex I impairment.

### Inhibition of lactate dehydrogenase impairs proliferation selectively in HTC cell lines

We hypothesized that the unique reliance on glycolysis for HTC cell survival predicted greater sensitivity to small molecule inhibitors of glycolytic enzymes compared to respiration competent cell lines. Thyroid cancer cell lines were exposed to 2-deoxyglucose (2-DG) and POMHEX, which inhibit hexokinase and enolase, respectively (Figure 3A) (Lin et al., 2020; Luengo et al., 2017; Wick et al., 1957). 2-DG and POMHEX treatment caused a dose-dependent decrease in proliferation and viability across all cell lines, with a small window of selectivity for complex I-mutant HTC (Figures 3B and 3C). Because glycolysis contributes to ATP production, we also assessed viability using an alternative, ATP-independent viability assay and obtained similar results (Figures S3A and S3B). Although 2-DG and POMHEX provided limited selectivity, these compounds displayed greater efficacy (reduction in cell viability at a compound’s maximum inhibitory concentration) against complex I-mutant HTC cells compared to respiration competent cell lines, which was consistent with the observed reliance on glycolysis for viability (Figure 2F).

**Figure 3:**
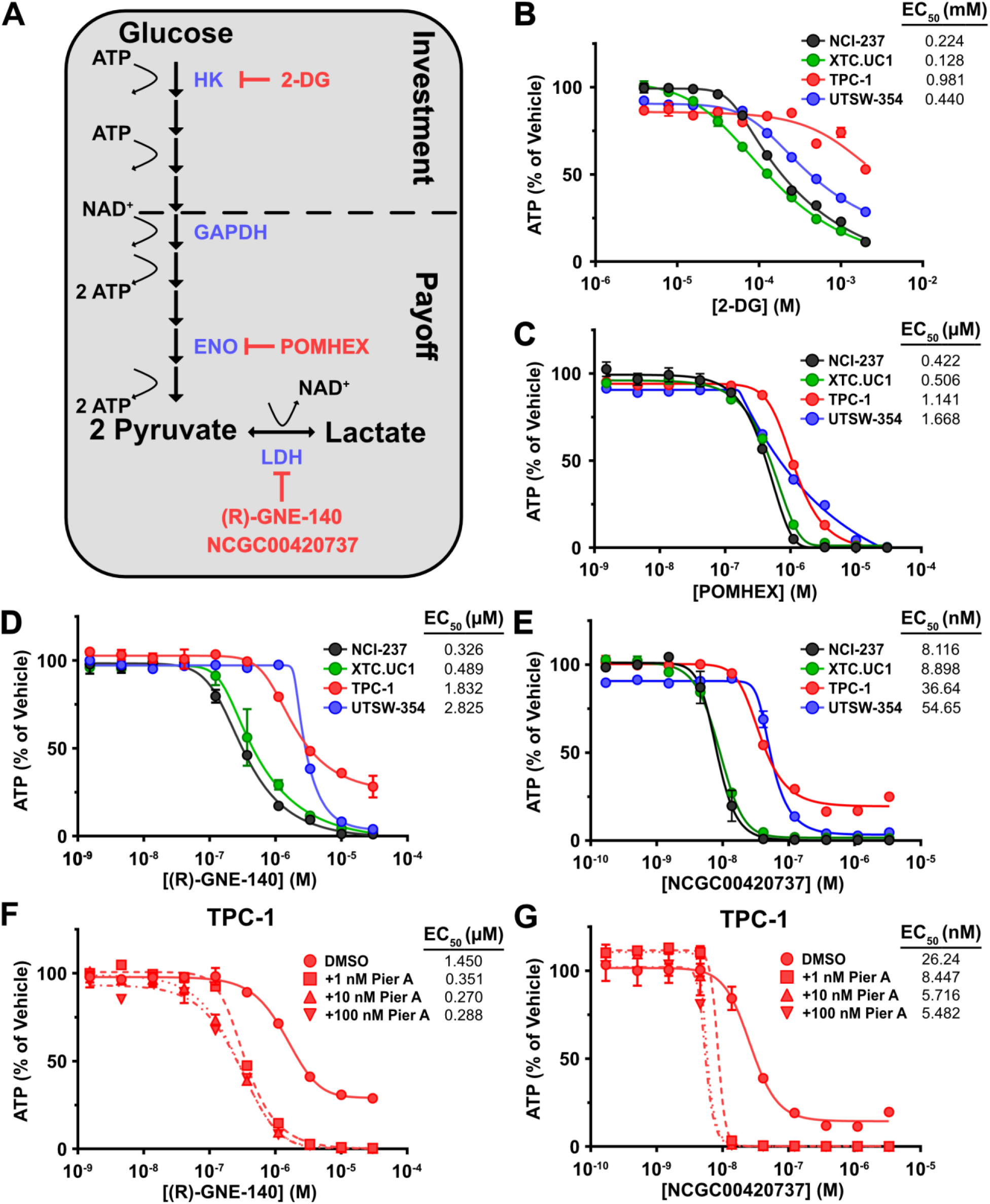
Inhibition of lactate dehydrogenase impairs proliferation selectively in HTC cell lines. A) Schematic depicting glycolysis and fermentation with inhibitors. Enzyme abbreviations are in blue; inhibitor names are in red. 2-DG = 2-deoxyglucose. B) Viability assay for cells treated with 2-DG for 3 days. C) Viability assay for cells treated with POMHEX for 3 days. D) Viability assay for cells treated with (R)-GNE-140 for 3 days. E) Viability assay for cells treated with NCGC00420737 for 3 days. F) Viability assay for TPC-1 cotreated with (R)-GNE-140 and indicated doses of piericidin A for 3 days. G) Viability assay for TPC-1 cotreated with NCGC00420737 and indicated doses of piericidin A for 3 days. Data are plotted as mean ± SEM of 2 replicates.

When oxygen is limiting or cells harbor oxidative phosphorylation defects, pyruvate is converted to lactate via the lactate dehydrogenase (LDH) reaction in a process known as fermentation. In the setting of respiratory deficiency, the LDH reaction becomes the primary source of NAD^+^ regeneration. NAD^+^ acts as an electron acceptor for numerous biosynthetic reactions, including the GAPDH reaction. Although LDH isoforms were not hits in our CRISPR/Cas9 screens, which was likely due to expression of functionally redundant paralogs, we hypothesized that LDH inhibition would serve as an ideal mechanism for indirectly inhibiting glycolysis in HTC cells because respiration-competent cells can sustain NAD^+^ regeneration through intact complex I function.

We treated thyroid cancer cell lines with two chemically distinct LDHA/LDHB inhibitors, (R)-GNE-140 and NCGC00420737 (Figure 3A) (Boudreau et al., 2016; Rai et al., 2017; Rai et al., 2020). Treatment with both LDH inhibitors caused a dose-dependent decrease in proliferation across all cells; however, HTC cells were approximately 5.7-fold more sensitive to (R)-GNE-140 and approximately 5.4-fold more sensitive to NCGC00420737 than complex I-competent thyroid cancer cells (Figures 3D, 3E, S3C, and S3D). LDH inhibitors also displayed greater efficacy in HTC cell lines, with NCGC00420737 causing an approximately 75% reduction in HTC cell viability at concentrations that had no effect on TPC-1 and UTSW-354 viability (Figures 3E and S3D). The selectivity and differences in efficacy we observed suggest that cells with sufficient respiratory capacity could survive acute LDH inhibition and provided a potential therapeutic window for LDH inhibitor application.

To test whether sensitivity to LDH inhibition was directly linked to complex I impairment, we treated complex I-competent cells with (R)-GNE-140 or NCGC00420737 and the complex I inhibitor piericidin A. Cotreatment with piericidin A caused a dose-dependent increase in sensitivity to LDH inhibitors in TPC-1 and UTSW-354 cells. We also noted a decrease in viability in piericidin-treated cell lines at higher concentrations, suggesting an induction of cell death similar to that observed in HTC cells (Figures 3F, 3G, S3E, and S3F). To exclude the possibility that LDH isoform expression could explain differences in sensitivity to LDH inhibition, we performed mRNA sequencing on NCI-237^UTSW^, TPC-1, and UTSW-354 cells. All three cell lines expressed similar levels of *LDHA* and *LDHB*, with minimal expression of *LDHC* and *LDHD* (Figure S3G). Taken together, these data suggest that complex I dysfunction was a key determinant of HTC sensitivity to LDH inhibition.

### ATP crisis and cell death follow LDH inhibition in complex-I mutant HTC cell lines

To understand the cause of impaired proliferation and cytotoxicity in HTC cells, we characterized the metabolic effects of LDH inhibition. As a major NADH-oxidizing center (Patgiri et al., 2020), we measured the NADH:NAD^+^ ratio in cells subjected to LDH inhibition. Treatment with NCGC00420737 or (R)-GNE-140 caused a dose-dependent increase in the cellular NADH:NAD^+^ ratio with similar EC_50_ values, suggesting target engagement at similar concentrations across all cell lines (Figures 4A and S4A).

**Figure 4:**
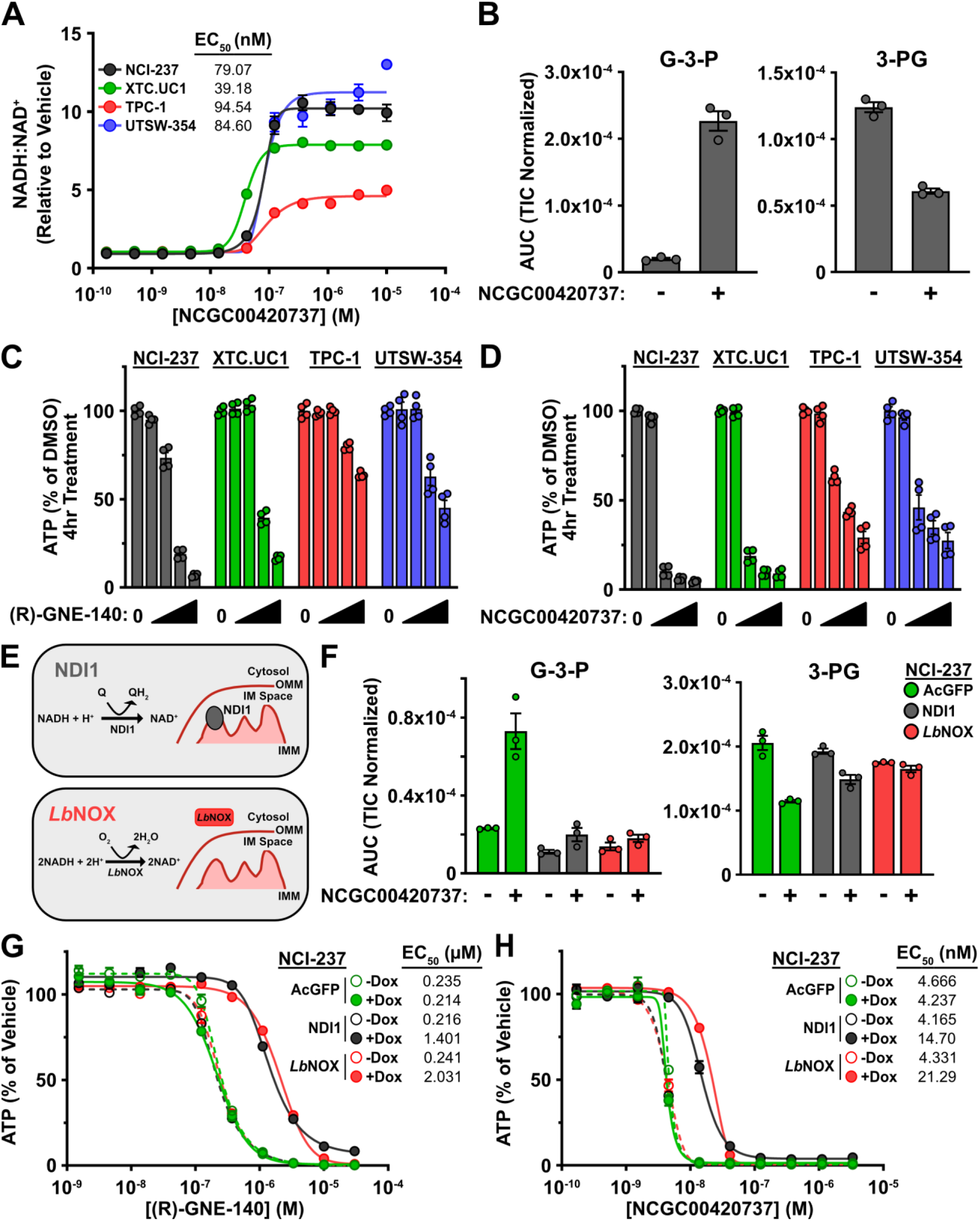
NAD^+^ regeneration rescues the effects of LDH inhibition. A) NADH:NAD^+^ ratio in cells treated with NCGC00420737 for 4 hours; *n* = 2 replicates. B) Metabolite levels in NCI-237^UTSW^ cells treated with 40 nM NCGC00420737 for 4 hours; *n* = 3 replicates. G-3-P = glyceraldehyde-3-phosphate; 3-PG = 3-phosphoglycerate. C) ATP levels in cells treated with (R)-GNE-140 for 4 hours; *n* = 4 replicates from 2 independent experiments. Concentrations are 0.041, 0.37, 3.33, and 30 µM (R)-GNE-140. D) ATP levels in cells treated with NCGC00420737 for 4 hours. Concentrations are 0.014, 0.123, 1.11, and 10 µM NCGC00420737. E) Schematic depicting reactions performed by NDI1 and *Lb*NOX, as well as cellular localization. OMM = outer mitochondrial membrane; IMM = inner mitochondrial membrane. F) Metabolite levels in NCI-237^UTSW^ cells expressing AcGFP, NDI1, or *Lb*NOX treated with 40 nM NCGC00420737 for 4 hours; *n* = 3 replicates. G-3-P = glyceraldehyde-3-phosphate; 3-PG = 3-phosphoglycerate. G) Viability assay for NCI-237^UTSW^ cells with (+Dox) or without (-Dox) expression of indicated construct treated with (R)-GNE-140 for 3 days; *n* = 2 replicates. H) Viability assay for NCI-237^UTSW^ cells with (+Dox) or without (-Dox) expression of indicated construct treated with NCGC00420737 for 3 days; *n* = 2 replicates. Data are plotted as mean ± SEM of indicated number of replicates.

We hypothesized that impaired NAD^+^ regeneration following LDH inhibition would inhibit glycolysis at GAPDH, which uses NAD^+^ as a cofactor. We used GC-MS to measure the levels of glycolytic intermediates in NCI-237^UTSW^ cells treated with NCGC00420737 or (R)-GNE-140. Consistent with GAPDH inhibition, we observed accumulation of the GAPDH substrate, glyceraldehyde-3-phosphate (G-3-P; 11.6- to 29.3-fold increase), and its isomer, dihydroxyacetone phosphate (DHAP; 6.5- to 7.9-fold increase), following LDH inhibitor treatment (Figures 4B, S4B, and S4C). In addition, we observed depletion of the lower glycolytic intermediate 3-phosphoglycerate (3-PG; approximately 2-fold decrease) (Figures 4B and S4C). Based on these data, we concluded that LDH inhibition blocked glycolytic flux in NCI-237^UTSW^ cells by inhibiting GAPDH.

The steps of glycolysis preceding GAPDH consume two molecules of ATP while the post-GAPDH steps yield four ATP molecules, for a net yield of two molecules of ATP per molecule of glucose. In the absence of oxidative phosphorylation, glycolysis becomes the primary ATP-producing center of the cell. We hypothesized that LDH inhibition and consequent disruption of glycolysis caused rapid ATP depletion in complex-I mutant HTC cells. We exposed cells to increasing concentrations of (R)-GNE-140 or NCGC00420737 for four hours and measured intracellular ATP levels. (R)-GNE-140 treatment caused a dose-dependent decrease in ATP levels for all cell lines, with HTC cells exhibiting the greatest decrease in ATP (1.6- to 2.2-fold decrease for TPC-1 and UTSW-354 and 5.9- to 14.6-fold decrease for NCI-237^UTSW^ and XTC.UC1) (Figure 4C). Treatment with NCGC00420737 had similar effects on cellular ATP (3.4- to 3.6-fold decrease for TPC-1 and UTSW-354 and 11.1- to 18.5-fold decrease for NCI-237^UTSW^ and XTC.UC1) (Figure 4D). As an orthogonal measure of ATP depletion, we assessed the phosphorylation of AMPK-activated protein kinase alpha (AMPKα) on Thr172, a well-defined signaling event that occurs following decreases in the cellular ATP:AMP ratio (Hawley et al., 1996; Stein et al., 2000; Trefts and Shaw, 2021). Consistent with decreases in cellular ATP, LDH inhibition caused a dose-dependent increase in AMPKα^T172^ phosphorylation in NCI-237^UTSW^ cells (Figure S4D). These data demonstrated that LDH inhibition induced rapid ATP depletion in complex I-mutant HTC, providing a mechanistic explanation for the cytotoxicity we observed uniquely in complex I-mutant HTC cells.

### NAD^+^ regeneration rescues the effects of LDH inhibition

If the impairment of GAPDH and resulting ATP crisis stemmed directly from the failure to regenerate NAD^+^ after LDH inhibition, then restoring NAD^+^ levels independent of LDH activity should completely rescue glycolysis, ATP levels, and cell viability. To accomplish this, we generated NCI-237^UTSW^ cells with lentiviral integration of doxycycline-inducible cassettes for expression of FLAG epitope-tagged AcGFP or two NADH-oxidizing enzyme systems with distinct mechanisms: NDI1, a yeast NADH dehydrogenase localized to mitochondria that catalyzes the transfer of electrons from NADH to ubiquinone yielding NAD^+^ and ubiquinol, and *Lb*NOX, a bacterial NADH oxidase localized to the cytosol that catalyzes the transfer of electrons from NADH to oxygen yielding NAD^+^ and water (Figure 4E) (Seo et al., 1998; Titov *et al*., 2016). Doxycycline-dependent expression of NDI1 and *Lb*NOX, but not AcGFP, increased the cellular NAD^+^:NADH ratio, suggesting these tools could be used to regenerate NAD^+^ following LDH inhibition (Figures S4E and S4F).

We investigated the effects on glycolysis following LDH inhibition when cells were provided an alternative mechanism for NAD^+^ regeneration. We induced AcGFP, NDI1, or *Lb*NOX expression in NCI- 237^UTSW^ cells and treated with NCGC00420737 or (R)-GNE-140. Treatment with either LDH inhibitor caused the accumulation of G-3-P and DHAP, and a corresponding decrease in 3-PG, in NCI-237^UTSW^ cells expressing AcGFP (Figures 4F, S4G, and S4H). Sustained NAD^+^ regeneration via expression of NDI1 or *Lb*NOX mitigated these effects, as indicated by similar levels of these glycolytic intermediates in both vehicle- and LDH inhibitor-treated cells (Figures 4F, S4G, and S4H). Additionally, expression of NDI1 or *Lb*NOX, but not AcGFP, sustained cellular ATP levels following short-term LDH inhibitor treatment (Figures S4I and S4J). Consistent with restoration of glycolytic activity and ATP levels, NDI1 and *Lb*NOX expression rescued NCI-237^UTSW^ proliferation following LDH inhibitor treatment, as indicated by an increase in EC_50_ (approximately 6.7- to 8.4-fold and 3.5- to 5.0-fold for (R)-GNE-140 and NCGC00420737, respectively), while expressing AcGFP had no effect (Figures 4G and 4H). Although NDI1 and *Lb*NOX expression caused similar EC_50_ increases, only NDI1 altered the efficacy of LDH inhibitor treatment, reflected by the upward shift in viability for NDI1-expressing cells at EC_100_ concentrations of LDHi (Figures 4G and 4H). Combined, the above data demonstrate that impaired NAD^+^ regeneration, inhibition of glycolysis at GAPDH, and ATP depletion underly cell death following LDH inhibition in HTC cells, rather than an unknown off-target effect of the small molecule inhibitors.

### LDH inhibition impairs HTC PDX Growth

We next tested whether LDH inhibition was efficacious against complex I-mutant HTC *in vivo*. NCI-237-R^PDX^ was implanted subcutaneously in immunodeficient mice and NCGC00420737 was administered at 30 mg/kg or 60 mg/kg intravenously for two weeks (two cycles of 5 days on, 2 days off). Treatment with 60 mg/kg NCGC00420737 impaired NCI-237-R^PDX^ growth and resulted in smaller tumors while administration of 30 mg/kg NCGC00420737 did not significantly affect NCI-237-R^PDX^ growth relative to vehicle (Figures 5A, 5B, and S5A). Animals receiving either dose of NCGC00420737 did not display signs of overt toxicity; however, all animals receiving compound had decreased red blood cell content and animals receiving 60 mg/kg NCGC00420737 exhibited weight loss that was recovered following compound withdrawal (Figures S5B and S5C).

**Figure 5:**
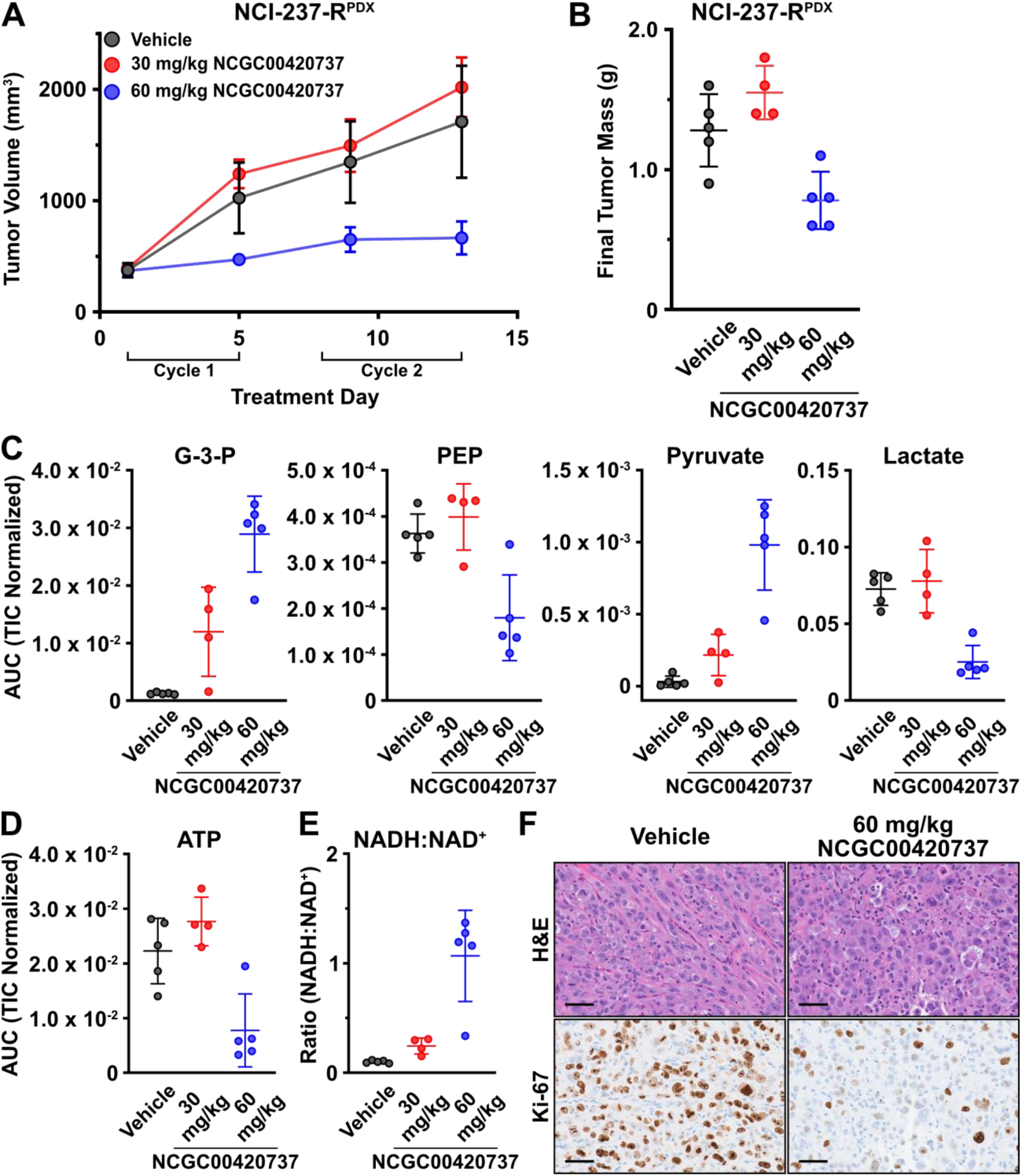
LDH inhibition impairs HTC PDX Growth. A) Tumor volumes for NCI-237-R^PDX^ xenografts treated with vehicle (*n* = 5), 30 mg/kg NCGC00420737 (*n* = 4), or 60 mg/kg NCGC00420737 (*n* = 5). Vehicle or compound was administered once daily via jugular vein catheter for two treatment cycles (5 days on, 2 days off). A final compound administration was performed on Day 6 of Cycle 2 and animals were sacrificed 1 hour after receiving compound. B) Final tumor mass for NCI-237-R^PDX^ xenografts from A). C) Metabolite levels in NCI-237-R^PDX^ xenografts from A) determined by LC-MS/MS. G-3-P = glyceraldehyde-3-phosphate; PEP = phosphoenolpyruvate. D) ATP levels in NCI-237-R^PDX^ xenografts from A) determined by LC-MS/MS. E) NADH:NAD^+^ ratio in NCI-237-R^PDX^ xenografts from A) determined by LC-MS/MS. F) Representative histology for NCI-237-R^PDX^ xenografts from A). Hematoxylin and eosin (H&E)-stained tumor sections (top). Immunohistochemical staining for Ki-67 in tumor sections (bottom). Scale bar represents 50 µm. Data are plotted as mean ± SD of indicated number of replicates.

To assess target engagement in tumors, we performed liquid chromatography-mass spectrometry (LC-MS/MS)-based metabolite profiling on tumors collected 1 hour after the final dose of vehicle or NCGC00420737 was administered. We observed a dose-dependent increase in intratumoral pyruvate and a decrease in lactate levels, consistent with target engagement and LDH inhibition (Figures 5C, S5D, and S5E). Additionally, we observed accumulation of upper (G-3-P) and depletion of lower (phosphoenolpyruvate, PEP) glycolytic intermediates, increases in the intratumoral NADH:NAD^+^ ratio, depletion of intratumoral ATP, and activation of AMPK signaling (Figures 5C-5E and S5F). These metabolic changes align with those observed upon LDH inhibition in cell culture (Figures 4A-4D and S4B-S4D). NCGC00420737 levels in plasma and tumors were higher in animals receiving 60 mg/kg, which likely explained the pronounced metabolic effects and tumor growth suppression observed in this treatment group (Figures S5D, S5E, and S5G).

Histological analysis of NCI-237-R^PDX^ tumors revealed a significant reduction in Ki-67 immunohistochemical staining in NCGC00420737-treated tumors, indicating a decrease in the number of actively proliferating cells (Figure 5F). Additionally, NCGC00420737-treated tumors displayed increased nuclear fragmentation relative to vehicle-treated tumors, suggesting the induction of cell death following LDH inhibition *in vivo* (Figure 5F). These data demonstrate that LDH inhibition was efficacious against complex I-mutant HTC *in vivo* and that a therapeutic window exists for targeting fermentation in pre-clinical HTC models with impaired complex I function.

## DISCUSSION

The selective pressure(s) that drive enrichment of damaging homoplasmic mtDNA mutations and impaired complex I activity in the setting of HTC and other cancers remain an open question. Here, we explore the metabolic consequences and therapeutic implications of this extreme enrichment for disruptive mtDNA mutations in HTC. Our results establish that near-homoplasmic damaging mtDNA mutations in HTC lead to ETC impairment and an obligate reliance on glycolysis for cell survival. HTC thus represents a true Warburg cancer in that damaged respiration underlies aerobic fermentation. The work presented here demonstrates that complex I function is dispensable for HTC tumor growth and that loss of mitochondrial respiration exposes LDH as a targetable metabolic liability in HTC.

How central carbon metabolism responds to respiratory defects in the context of cellular proliferation has largely been informed by studies of cytoplasmic hybrid (cybrid) and cultured cells subjected to pharmacologic inhibition of the ETC (Birsoy *et al*., 2015; Mullen *et al*., 2011; Sullivan *et al*., 2015). Whether these adaptations occur in the context of naturally arising complex I loss, as exemplified by HTC, was heretofore unexplored. We employed pooled genome-wide and targeted CRISPR/Cas9 screens to systematically profile essential genes in HTC cells, and our screens identified glycolysis as a selective vulnerability in complex I-mutant HTC cells despite the fact that 6 out of 10 glycolytic enzymes are reported as ‘Common Essential’ in The Cancer Dependency Map (www.depmap.org/portal/). This concept draws support from previous genetic screens that identified similar relationships wherein depletion of mitochondrial ETC components was a liability for cells grown under limiting glucose conditions (Arroyo *et al*., 2016; Birsoy et al., 2014). Additionally, cancer cell lines harboring heteroplasmic mtDNA mutations in complex I subunits could not upregulate oxidative phosphorylation to a sufficient level to maintain proliferation under limiting glucose conditions (Birsoy *et al*., 2014). These results highlight the power of genetic screens to unveil selectivity in essential cellular processes and suggest a therapeutic window for targeting glycolysis in complex I-mutant HTC.

Considering the universal essentiality of glycolytic enzymes, targeting glycolysis by direct inhibition of these enzymes presents a major pharmacologic challenge. Precise titration of inhibitors would be required to achieve a therapeutic benefit without systemic effects; however, such precision is likely unattainable for most cancers. While isoform-specific inhibitors or isoform-specific expression of enzymes could afford a therapeutic window, these cases are also likely to be infrequent (Lin *et al*., 2020). Illustrating the difficulty in directly targeting glycolysis, we observed that 2-DG and POMHEX provided negligible selectivity for HTC cells. These results raise the question of why glycolytic enzymes scored as HTC-selective in our CRISPR/Cas9 screens. When cells were cultured in glucose-free or galactose-containing media, we observed a marked reduction in proliferation across all cell lines, yet we only observed decreases in viability for complex I-mutant HTC cells. Similarly, 2-DG and POMHEX impaired proliferation irrespective of complex I status; however, we observed greater decreases in cell viability in complex I-mutant HTC cells.

Complex I is a major NAD^+^ regenerating center and loss of ETC function shifts cellular NAD^+^ regeneration to LDH. The longstanding observation that cells with ETC defects require exogenous pyruvate for cell growth reinforces the importance of fermentation in the setting of respiratory dysfunction as this pyruvate is used to drive NAD^+^ regeneration via LDH (King and Attardi, 1989; Sullivan *et al*., 2015; Titov *et al*., 2016). The defects in NAD^+^ regeneration exposed by complex I loss in HTC manifest in the form of pyruvate auxotrophy and increased sensitivity to LDH inhibition. We demonstrated that targeting LDH pharmacologically disrupts cellular redox and impairs glycolysis at GAPDH, which requires NAD^+^. The rescue of glycolytic flux and cell viability following expression of *Lb*NOX or NDI1, which regenerate NAD^+^ independently of LDH, reinforced that these effects were due to on-target LDH inhibition. Therefore, intact complex I, and more broadly, respiratory activity buffer against LDH inhibition by providing an additional mechanism for NAD^+^ regeneration.

While impaired NAD^+^ regeneration has myriad consequences for anabolic metabolism and cell proliferation, we speculate that disruption of ATP synthesis is the underlying cause of cytotoxicity following LDH inhibition in HTC cells. Inhibiting glycolysis at GAPDH decouples the ‘investment’ and ‘payoff’ phases of glycolysis. As a result, LDH inhibition transforms the primary ATP-generating pathway in cells with ETC dysfunction into an ATP-consuming process. Consistent with this concept, we observed rapid ATP depletion and cell death in HTC cells following LDH inhibition. Additional support for this assertion comes from our observation that culturing cells in media with galactose as a primary carbon source, which results in inefficient glycolytic ATP synthesis, results in a marked decline in HTC cell viability. By contrast, pyruvate withdrawal, which impairs NAD^+^ regeneration but does not have substantial effects on ATP synthesis, impaired complex I-mutant HTC cell growth but had no effects on cell viability. In addition, expression of NDI1, which re-engages the mitochondrial ETC and supports limited ATP production through oxidative phosphorylation, partially rescued cell death in response to LDH inhibition. An impaired capacity for oxidative phosphorylation thus underlies the cytotoxicity observed in complex I-mutant HTC and exposes a therapeutic window for LDH inhibition in HTC.

We demonstrate that the direct effects of LDH inhibition observed in cultured cells translate to the PDX model of HTC. Two cycles of treatment with 60 mg/kg NCGC00420737 induced a marked NCI-237-R^PDX^ tumor growth response with limited adverse effects, providing preclinical evidence that LDH is a tractable target in complex I-mutant HTC. Metabolite profiling in NCI-237-R^PDX^ tumors treated with NCGC00420737 reinforce that these effects were due to on-target LDH inhibition as we observed metabolic changes consistent with those observed in culture. Similar to previous reports (Yeung et al., 2019), we observed signs of hemolysis following NCGC00420737 administration. The observance of hemolysis, and descriptions of aerobic glycolysis in normal tissues including the pigmented retinal epithelium and lymphocytes, necessitate close monitoring of these cells and tissues during continued preclinical evaluation of LDH inhibitors. These results, however, provide encouraging evidence that glucose metabolism represents a tractable target in complex I-mutant HTC.

Mitochondrial accumulation and evidence of metabolic rewiring are defining features of HTC, yet approximately 40% of HTC tumors do not harbor discrete mutations in complex I subunits (Ganly et al., 2022; Ganly *et al*., 2018; Ganly and McFadden, 2019; Gopal *et al*., 2018; McFadden and Sadow, 2021). This raises the question whether all HTC tumors exhibit functional ETC impairment. We observed that XTC.UC1 cells harbor a heteroplasmic loss-of-function mutation in *MT-ND1*, and although respiration was depressed relative to cell lines lacking homoplasmic loss-of-function mtDNA mutations, these cells still displayed ETC-linked oxygen consumption (Figure 1C). Nevertheless, culture of XTC.UC1 in glucose-free or galactose-containing media suppressed viability, and XTC.UC1 cells were sensitive to LDH inhibition. These results suggest that ETC function was below a threshold required for viability following LDH inhibition or culture in glucose-free or galactose-containing media. Analysis of glucose metabolism in XTC.UC1 revealed that these cells display defects in glucose oxidation and recent profiling of patient-derived HTC tumors identified metabolic changes consistent with ETC dysfunction (Ganly *et al*., 2022). Therefore, HTC tumors that lack damaging near-homoplasmic mtDNA mutations might harbor sufficient disruption of oxidative phosphorylation and glucose metabolism to be sensitized to LDH inhibition. While the widespread application of LDH inhibitors in HTC requires additional investigation, results from this study and other studies that utilized a combination small molecule approach reinforce the synthetic lethal interaction between complex I dysfunction and LDH inhibition (Oshima et al., 2020). Our study raises the possibility that near-homoplasmic mtDNA mutation status could be used as a biomarker for guiding LDH inhibitor-based therapeutic strategies in HTC. The continued development of HTC models and characterization of mitochondrial ETC function in HTC tumors will be important for a more complete understanding of HTC-specific metabolic alterations and sensitivity to LDH inhibition.

Despite the longstanding observation that tumors exhibit altered glucose metabolism, attempts to therapeutically target glucose metabolism in cancer have had limited success (Luengo *et al*., 2017; Warburg, 1956). Fermentation has been an attractive target in cancer, yet recent studies have highlighted the adaptive nature of central carbon metabolism that allows cells to respond to LDH inhibition by upregulating oxidative phosphorylation (Boudreau *et al*., 2016). Given the genetically defined and irreversible nature of mitochondrial ETC dysfunction in HTC, it is unlikely that HTC tumors challenged with LDH inhibition would be able to upregulate oxidative phosphorylation. Beyond HTC, the presence of high allelic fraction disruptive mtDNA mutations in colon and renal carcinomas suggests a potential therapeutic opportunity for LDH inhibition in these cancers (Yuan et al., 2020). Studies conducted in patients with renal cell carcinomas (RCC), as well as FH-deficient RCC models, demonstrate that these tumors display ETC dysfunction and alterations in glucose metabolism that could expose a tractable sensitivity to LDH inhibition (Courtney et al., 2018; Crooks et al., 2021). Thus, we propose extending preclinical studies with LDH inhibitors into other cancer types with genetically defined ETC lesions, or evidence of respiratory dysfunction, as a mechanism for identifying relevant therapeutic footholds in glucose metabolism.

## ACKNOWLEDGMENTS

The authors thank Florentina Moreno for technical and administrative support. We thank Alex Sternisha for assistance with GC-MS data analysis. We acknowledge Cheryl Lewis and the Simmons Comprehensive Cancer Center Tissue Management Shared Resource, which is supported by the National Cancer Institute, National Institutes of Health under award number P30 CA142543, for IHC assistance and slide imaging; John Shelton and the UT Southwestern Histo Pathology Core for histology assistance; Vanessa Schmid and the Eugene McDermott Center for Human Growth and Development Next Generation Sequencing Core for CRISPR/Cas9 screen sequencing; Hamid Baniasadi and the UT Southwestern Department of Biochemistry Metabolomics Core Facility for LC-MS/MS metabolite profiling; the Children’s Research Institute Metabolomics Facility for GC-MS metabolite profiling; and effort and resources provided by the institutionally supported UT Southwestern Preclinical Pharmacology Core. D.G.M. was supported by the Cancer Prevention and Research Institute of Texas (RR140884), a Disease-Oriented Scholar Award from UT Southwestern Medical Center, and the Damon Runyon Cancer Research Foundation (102-19). A.R.F. was supported by the NIH Pharmacological Sciences Training Grant (T32-GM007062). S.D.S. was supported by the National Science Foundation Graduate Research Fellowship (2019281210). P.M. was supported by the Cancer Prevention and Research Institute of Texas (RP180778). NCGC00420737 was developed using funding from the National Cancer Institute, National Institutes of Health under contract HHSN261200800001E, and with funding from the Chemical Biology Consortium, National Cancer Institute Experimental Therapeutics (NExT) Program. We thank members of the McFadden lab for helpful discussions and review of the manuscript and Deepak Nijhawan, Ben Tu, and Ralph DeBerardinis for critical review of the manuscript.

## AUTHOR CONTRIBUTIONS

D.G.M. conceived the study and supervised the work. A.R.F. and D.G.M. designed and interpreted experiments and A.R.F. performed experiments. V.L. performed experiments. S.D.S. and P.M. performed targeted mtDNA sequencing and analyzed mtDNA sequencing data. J.K. analyzed whole-exome sequencing, RNA-sequencing, and CRISPR/Cas9 screening datasets, supervised by Y.X. G.M.S. and L.N. provided NCGC00420737, as well as pharmacokinetic data and discussions related to NCGC00420737 administration in animals. N.W. assisted with animal studies and performed pharmacokinetic profiling of NCGC00420737 in animals. D.G.M. and A.R.F. wrote the manuscript.

## DECLARATION OF INTERESTS

The authors declare no competing interests.

## MATERIALS AND METHODS

### Patient-derived materials and human cell lines

The HTC PDX model, 248138-237-R [Lot# GF9] (male), was developed by the NCI PDMR. NCI-248138-237-R^PDX^ was cryorecovered and expanded in NOD.Cg-*Prkdc*^*scid*^ *Il2rg*^*tm1Wjl*^/SzJ mice (NSG™; Jackson Laboratories strain code 005557) prior to use. UTSW-354 (female) was derived from a papillary thyroid cancer metastasis to lymph node and obtained under IRB Protocols STU 072017-103 (PI – McFadden) and STU 102010-051 (PI – Lewis). A complete description of UTSW-354 will be reported elsewhere. NCI-237^UTSW^ cell line was generated by dissociating a freshly harvested NCI-237-R^PDX^ tumor using the Miltenyi MACS Human Tumor Dissociation Kit (Miltenyi 130-095-929). Tumor bearing animals were sacrificed by CO_2_ euthanasia, 1 NCI-237-R^PDX^ tumor (approximately 8 mm x 8 mm x 8 mm) was dissected, and approximately half of the tumor was minced using a sterile razor blade. Minced tissue was placed in 5 mL RPMI (Sigma R8758) containing 200 µL Enzyme H, 100 µL Enzyme R, and 25 µL Enzyme A that were prepared according to the manufacturer’s instruction and incubated for approximately 30 minutes at 37°C with agitation every 5-10 minutes. The resulting cell suspension was filtered through a 40 µm nylon mesh filter, pelleted, and resuspended in RPMI supplemented with 2% FBS (Sigma F0926), 2 mM L-glutamine (Sigma G7513), 1% penicillin/streptomycin (Sigma P4333), 1X MEM non-essential amino acids (Sigma M7145), 100 µM sodium pyruvate (Sigma S8636), 50 µg/mL uridine (Sigma U3003), and 1X insulin-transferrin-selenium (Sigma I1884). Cells were cultured using standard trypsinization and passaging techniques. UTSW-354 cell line was generated using a similar protocol. TPC-1 (female) was obtained from Dr. Sareh Parangi (Massachusetts General Hospital, Boston, MA, USA) and XTC.UC1 (female) was a gift from Dr. Ian Ganly (Memorial Sloan Kettering Cancer Center, New York, NY, USA). HEK293T/17 (female) were purchased from ATCC (CRL-11268). All cell lines were subjected to short-tandem repeat (STR) profiling performed by the Eugene McDermott Center for Human Growth and Development Sequencing Core. Cell lines were periodically tested for mycoplasma contamination using a PCR-based assay.

### Animal studies

Animal studies were approved by the UT Southwestern Institutional Animal Care and Use Committee. For NCI-237-R^PDX^ implantation, immunodeficient mice were anesthetized using isoflurane/oxygen. Tumor fragments were placed in the subcutaneous space, and incisions were closed using Vetbond (3M). For NCI-237 cell line xenograft initation, 2 × 10^6^ cells were injected subcutaneously (resuspended in 1:1 media:Geltrex™ (Thermo Fisher A1413201)).

### Cell culture

Cell lines were maintained at 37°C with 5% CO_2_. HEK 293T/17 were cultured in DMEM (Sigma D6429) supplemented with 10% FBS, 2 mM L-glutamine, 1% penicillin/streptomycin, and 50 µg/mL uridine. All experiments with thyroid cancer cell lines were performed in human plasma-like media (HPLM) unless otherwise indicated. Cell lines were adapted to culture in HPLM supplemented with 2% FBS, 1% penicillin/streptomycin, and 1X insulin-transferrin-selenium for at least 5 passages. HPLM pool stocks were prepared and stored as previously described (Cantor *et al*., 2017) and HPLM was assembled by combining pool stock solutions in deionized water, adjusting pH to ∼7.2 with hydrochloric acid, and filtering through a 0.22 µm PES filter (Corning 431097). Prepared HPLM was used within 3-4 days and supplements were added daily before use. For cell growth assays requiring pyruvate-free media, a separate ‘Pool 9 stock’ solution that lacked pyruvate was prepared and used to prepare HPLM -pyruvate before pH adjustment and sterile filtration. For cell growth assays requiring glucose-free media, glucose was omitted to prepare HPLM -glucose before pH adjustment and sterile filtration.

### NCGC00420737 animal dosing studies

8 week old male NOD-*Prkdc*^*em26Cd52*^*Il2rg*^*em26Cd22*^/NjuCrl (NCG; strain code 572) mice with jugular vein catheters (surgical code: JUGVEIN) and transcutaneous one-channel vascular access buttons (accessory code: INSTBUTON1CH) were purchased from Charles River Laboratories and individually housed at UT Southwestern Medical Center. NCI-237-R^PDX^ was cryorecovered and propagated in NOD.Cg-*Prkdc*^*scid*^ *Il2rg*^*tm1Wjl*^/SzJ mice (NSG™; Jackson Laboratories strain code 005557) for 4 weeks before subcutaneous implantation in NCG mice with jugular vein catheters. Tumors were measured weekly using calipers and tumor volume was calculated according to [volume = ((Length)*(Width)^2^)*3.14)/6]. Dosing was initiated when tumors reached approximately 300-500 mm^3^. NCGC00420737 was prepared as a 10 mg/mL stock solution that was dissolved in 0.1 N NaOH/PBS and pH was adjusted to approximately 7.4 using 1 N HCl. Vehicle was prepared using the same method. A fresh stock of NCGC00420737 or vehicle was prepared weekly and kept at 4°C, protected from light. NCGC00420737 was administered via jugular catheter using the transcutaneous vascular access button for 2 cycles of 5 days on, 2 days off. Tumor and weight measurements were performed daily. A final dose of compound was administered on Day 6 of Cycle 2 and animals were sacrificed by CO_2_ asphyxiation approximately 1-1.5 hours after receiving a final dose of NCGC00420737 or vehicle. Blood was collected from the inferior vena cava and immediately placed in potassium-EDTA tubes. Tumors were dissected, weighed, and snap-frozen in liquid nitrogen. One animal from the 60 mg/kg dosing group was taken off study after a single dose of compound due to signs of physical discomfort.

### Complete blood counts

Complete blood counts (CBCs) were performed by the UT Southwestern Medical Center Animal Resource Center on potassium-EDTA-treated whole blood that was stored at 4°C for approximately 24 hours prior to analysis.

### Respiration analysis

An Agilent Seahorse XFe96 Analyzer was used for respiration analysis. For intact cell respiration measurements, cells were plated at 20,000 cells/well in 80 µL media, allowed to adhere for 3 hours, supplemented with an additional 200 µL media, and cultured overnight. The following day, cells were rinsed twice using 200 µL/well Seahorse assay medium (DMEM (Sigma D5030) with 10 mM glucose, 2 mM L-glutamine, 1 mM sodium pyruvate, and 1% penicillin/streptomycin), and 150 µL assay medium was added after the second wash. Cells were allowed to equilibrate in a 37°C, CO_2_-free incubator for 45 minutes before assaying. Standard calibration was performed, and oxygen consumption measurements were performed using a 3 minute ‘Mix,’ 3 minute ‘Measure’ cycle. 3 measurements were collected at baseline and after the injection of each compound. Compounds were used at the following concentrations: 2 µM oligomycin, 3 µM CCCP, 1 µM piericidin A, and 3 µM antimycin A. For permeabilized cell respiration measurements, cells cultured in RPMI (media as described above) were plated at 20,000 cells/well in 80 µL media and allowed to adhere overnight. The following day, cells were rinsed twice using 200 µL/well Seahorse assay medium (MAS buffer containing 5 mM pyruvate, 2.5 mM malate, and 2 mM adenosine diphosphate (ADP) for complex I measurement or 10 mM succinate and 2 mM ADP for complex II measurement), and 150 µL respective assay medium containing 2 nM Seahorse XF Plasma Membrane Permeabilizer (Agilent 102504-100) was added after the second wash. Oxygen consumption measurements were immediately performed without standard calibration.

### Cell proliferation assays

Cells were plated in 6 well plates (Corning) in 2 mL media at the following densities: NCI-237 (30,000 cells/well); XTC.UC1 (30,000 cells/well); TPC-1 (30,000 cells/well); UTSW-354 (100,000 cells/well) and allowed to adhere overnight. The following day, cells were washed once with 2 mL PBS (Sigma D8537) and 8 mL of growth media was added to the respective wells. Growth media lacking pyruvate or glucose was prepared as described above and supplemented with 2% dialyzed FBS, 1% penicillin/streptomycin, and 1X insulin-transferrin-selenium. For pyruvate auxotrophy experiments, media was supplemented with 100 µM sodium pyruvate from a sterile 100 mM stock. For glucose withdrawal experiments, media was supplemented with 5 mM galactose or 5 mM glucose from sterile 500 mM stocks. Cells were cultured for the indicated number of days before trypsinization and counting on a ViCell™-XR Cell Viability Analyzer (Beckman Coulter). Fold-change in cell number was calculated relative to a baseline cell count obtained by counting cells from replicate plates that were collected on the day of media exchange. Growth conditions were assayed in triplicate and experiments were performed at least twice.

### Cell viability assays

Cells were plated in 6 well plates (Corning) in 2 mL media at 100,000 cells/well and allowed to adhere overnight. The following day, cells were washed once with 2 mL PBS and 8 mL of growth media was added to the respective wells. Growth media lacking pyruvate or glucose was prepared as described above and supplemented with 2% dialyzed FBS, 1% penicillin/streptomycin, and 1X insulin-transferrin-selenium. For pyruvate viability experiments, media was supplemented with 100 µM sodium pyruvate from a sterile 100 mM stock. For glucose viability experiments, media was supplemented with 5 mM galactose or 5 mM glucose from sterile 500 mM stocks. Cells were cultured for 48 hours before all cells (floating and adherent) were collected by trypsinization and pelleting at 500x *g* for 5 minutes. Cells were resuspended in 500 µL PBS + 0.5 µg/mL propidium iodide (Life Technologies P1304MP) and stained in the dark at room temperature for 10 min before data acquisition and data analysis on a Guava easyCyte™ flow cytometer (EMD Millipore). Conditions were assayed in triplicate and experiments were performed at least twice.

### Compound dose-response curves

Thyroid cancer cell lines were plated in 96 well plates (Corning 3903) in 100 µL media at the following densities: NCI-237 (2,000 cells/well); XTC.UC1 (2,000 cells/well); TPC-1 (1,000 cells/well); UTSW-354 (5,000 cells/well) and allowed to adhere overnight. The following day, the media was exchanged and 200 µL fresh media was added to each well. Compounds (POMHEX (MedChemExpress HY-131904), (R)-GNE-140 (MedChemExpress HY-100742A), and NCGC00420737 (kindly provided by G.M.S., L.N., and the NCI Experimental Therapeutics Program)) were dispensed in a 3-fold dilution series using a D300e Digital Dispenser (Tecan). Piericidin A (Cayman 15379) cotreatment was also performed using a D300e Digital Dispenser. 2-DG (Sigma D8375) was administered in a 2-fold dilution series that was performed by hand using a multi-channel pipette. Cells were cultured with compound for 72 hours before viability was assessed using an ATP-based luminescent viability assay, CellTiter-Glo (Promega G7570). Plates were cooled to room temperature for 30 minutes, 100 µL growth media was removed, and 40 µL CellTiter-Glo (prepared according to manufacturer’s directions and diluted 1:1 with PBS + 1% Triton-X-100) was added to each well. Plates were incubated for 10 minutes at room temperature, shaking at 120 rpm on an orbital shaker, before data acquisition using a Synergy 2 or Cytation 5 plate reader (BioTek). For ATP-independent viability assays, cells were cultured with compounds as above before incubation with glycylphenylalanyl-aminofluorocoumarin (GF-AFC). Cells were incubated with 100 µM GF-AFC for 3 hours at 37°C in a 5% CO_2_ incubator, plates were cooled at room temperature for 30 minutes in the dark, and data was acquired using a Cytation 5 plate reader (BioTek) using 400 nm excitation and 520 nm emission filters. GF-AFC was provided by Dr. Deepak Nijhawan (UT Southwestern Medical Center, Dallas, TX, USA). Background fluorescence values (media only + GF-AFC) were subtracted before data analysis. For all dose-response curves, data were normalized relative to vehicle-only wells and EC_50_ values were determined using Prism (GraphPad), asymmetric (five parameter), least squares curve fitting. Conditions were assayed in duplicate and all experiments were performed at least twice.

### NADH/NAD^+^ measurements

Thyroid cancer cell lines were plated in 96 well plates (Corning 3903) in 100 µL media at the following densities: NCI-237 (10,000 cells/well); XTC.UC1 (20,000 cells/well); TPC-1 (20,000 cells/well); UTSW-354 (30,000 cells/well) and allowed to adhere overnight. The following day, the media was exchanged and 200 µL fresh media was added to each well. Compounds were dispensed in a 3-fold dilution series by hand using a multi-channel pipette. Cells were cultured for 4 hours at 37°C in a 5% CO_2_ incubator before samples were collected for metabolite analysis. Media was quickly removed using a multi-channel pipette and cells were lysed with 100 µL/well ice cold lysis buffer (equal parts 1% DTAB/0.2N NaOH and PBS) and immediately placed at −80°C. Samples were thawed on ice and divided into 96 well PCR plates (40 µL lysate/well). For NADH measurement, the plate was sealed, incubated at 60°C for 15 minutes on a thermal cycler, cooled to room temperature for 20 minutes, and 40 µL 1:1 0.4 N HCl:0.5 M Tris base was added to each well, followed by thorough mixing. For NAD^+^ measurement, 20 µL 0.4 N HCl was added to each well, mixed thoroughly, the plate was sealed, incubated at 60°C for 15 minutes on a thermal cycler, cooled to room temperature for 20 minutes, and 20 µL 0.5 M Tris base was added to each well, followed by thorough mixing. NAD/NADH-Glo (Promega G9071) detection reagent was prepared according to the manufacturer’s instructions, 10 µL detection reagent was mixed with 10 µL sample in a white-walled 384 well plate and incubated for 1 hour at room temperature before data acquisition using a Synergy 2 plate reader (BioTek). Individual readings for NADH and NAD^+^ were compared to generate NADH:NAD^+^ ratios, data were normalized relative to vehicle-only wells, and EC_50_ values were determined using Prism (GraphPad), asymmetric (five parameter), least squares curve fitting. The same protocol was used for determining baseline NADH:NAD^+^ ratios in NCI-237 pZIP expressing AcGFP, NDI1, or *Lb*NOX.

### ATP measurements

Cells were plated and compound was administered as above for NADH/NAD^+^ measurements. After 4 hour incubation, plates were cooled to room temperature for 30 minutes, 100 µL growth media was removed, and 40 µL CellTiter-Glo (prepared according to manufacturer’s directions and diluted 1:1 with PBS + 1% Triton-X-100) was added to each well. Plates were incubated for 10min at room temperature, shaking at 120 rpm on an orbital shaker, before data acquisition using a Synergy 2 or Cytation 5 plate reader (BioTek). Data were normalized to vehicle-only wells.

### Lentivirus production

All lentiviruses were produced by co-transfection of HEK 293T/17 cells with plasmid DNA for the respective lentiviral vector and the packaging components psPAX2 and pMD2.G at a 5:3:2 mass ratio (vector:psPAX2:pMD2.G) using TransIT-LT1 (Mirus MIR 2300). Lentiviral supernatants were collected at 48 and 72 hours post-transfection, pooled, and filtered using 0.45 µm PES filters before immediate use or aliquoting and storage at −80°C. psPAX2 and pMD2.G are gifts from Didier Trono (Addgene #12260 and 12259, respectively).

### Generating NCI-237 AcGFP, NDI1, and *Lb*NOX cells

Plasmids containing AcGFP and LbNOX (Addgene #75285) were gifts from Dr. Deepak Nijhawan (UT Southwestern Medical Center, Dallas, TX, USA) and Dr. Vamsi Mootha (Massachusetts General Hospital, Boston, MA, USA), respectively. Human codon optimized NDI1 was based on the sequence described in (Titov *et al*., 2016) and purchased as a gBlock from IDT. Lentiviral vectors for doxycycline-inducible expression of AcGFP, NDI1, and *Lb*NOX were generated by PCR amplification of the respective cDNAs and Gibson assembly of cDNA fragments and digested vector. Lentiviruses were produced as described above and NCI-237 cells were transduced with increasing volumes of lentiviral supernatant. Forty-eight (48) hours after transduction, cells were collected and plated directly into media containing 2 µg/mL puromycin and cultured for 48 hours. Cells transduced at a high MOI (∼90-95% cell survival based on visual estimate) were selected and expanded. For all experiments requiring AcGFP, NDI1, and *Lb*NOX expression, cells were plated directly into media containing 100 ng/mL doxycycline and cultured for at least 16 hours before starting treatment or collecting samples for analysis.

### Genome-wide CRISPR/Cas9 screening

NCI-237 and TPC-1 cells line stably expressing Cas9 were generated by infecting cells with lentiCas9-Blast at an MOI of ∼0.5 (visual estimation) (lentiCas9-blast is a gift from Feng Zhang, Addgene # 52962) and selecting cells with 10 µg/mL blasticidin (Invivogen ant-bl-1). The human Brunello CRISPR knockout pooled sgRNA library was a gift from David Root and John Doench (Addgene #73178). sgRNA library lentivirus was produced via large-scale transfection of 20 × 150mm dishes of HEK293T/17 (12 × 10^6^ cells/dish were plated and transfected with 30 µg plasmid DNA the following day). Lentiviral titers were assessed by determining the fraction of puromycin-resistant cells following lentivirus exposure. Cas9-expressing cells were exposed to increasing amounts of lentiviral supernatant for approximately 18 hours, media was changed, and cells were cultured for an additional 30 hours. 48 hours after exposure to virus, cells were collected, counted, and equally divided into media with or without 2 µg/mL puromycin (Sigma P8833). Cells were cultured for 48 hours in puromycin-containing media and the fraction of puromycin-resistant cells was calculated by comparing the number of cells in media +puromycin to the number of cells in media-puromycin. Viral titers that gave 20-40% cell survival were used for large-scale infection and screening. For screening, cells were cultured in RPMI-based media as described above and an initial pool of Cas9-expressing cells (approximately 270 × 10^6^ cells per cell line) were infected to achieve at least 1,000X cell coverage of the sgRNA library after puromycin selection. Puromycin selection was initiated 48 hours after exposure to virus, and cells were cultured in puromycin-containing media for 7 days before an initial cell population (approximately 300 × 10^6^ cells) was collected. Approximately 80 × 10^6^ cells were maintained at each passage and cells were grown for approximately 12 population doublings before a final cell population was collected. Genomic DNA was prepared using the Blood and Cell Culture DNA Maxi Kit (Qiagen 13362). To determine sgRNA distribution in the initial and final cell populations, sgRNA cassettes were PCR-amplified to attach adapters and barcode sequences for next-generation sequencing. sgRNA cassettes were amplified using 22 cycles of PCR amplification from approximately 300 µg genomic DNA (10 µg DNA/100 µL reaction) using 2X KAPA HiFi HotStart ReadyMix (Roche KK2602), a combination of 8 staggered P5 primers, and 1 unique P7 index primer per sample as described by the Broad Institute Genetic Perturbation Platform (www.portals.broadinstitute.org/gpp/public/resources/protocols). Individual reactions for each sample were pooled and purified using AMPure XP bead-based purification (Beckman Coulter A63881) before sequencing on an Illumina NextSeq 500 with 75 duty cycles. BWA was used to map the sequencing reads to the sgRNA sequences concatenated with 5’ and 3’ flanking sequences. The mapping reads were counted for each sgRNA using SAMtools (v1.9) (Li et al., 2009) with the options “-F 2308 -q 1”. MAGeCK (Li et al., 2014) was used to determine the relative depletion and enrichment of genes in treatment samples compared to the control samples.

### Targeted CRISPR/Cas9 screening

Genes were selected for targeted CRISPR/Cas9 screening based on the following criteria: i) the gene displayed ≥ 2-fold dropout in NCI-237 cells in the genome-wide screen; ii) the gene displayed ≥ 1.5-fold greater dropout in NCI-237 than TPC-1 in the genome-wide screen; iii) the gene was not depleted in > 99% of cell lines in The Cancer Dependency Map (www.depmap.org/portal/; DepMap Public 21Q2 Assembly) The targeted validation library sgRNA sequences were obtained from the Brunello library and synthesized as an oligo pool containing the sgRNA sequences with overhangs for Gibson assembly (Twist Biosciences). The sgRNA plasmid DNA library was constructed as described in (Joung et al., 2017) and sgRNA distribution in the library was assessed by next-generation sequencing. Cell lines conditioned to culture in HPLM or RPMI were infected with lentiCas9-Blast to generate cell lines stably expressing Cas9. Lentiviral titers were determined as for the genome-wide CRISPR/Cas9 screens and cell lines were infected in duplicate (approximately 8 × 10^6^ cells per replicate) to achieve at least 1,000X cell coverage of the sgRNA library after puromycin selection. Cells were selected with puromycin as above before collecting an initial cell population (approximately 8 × 10^6^ cells per replicate). Approximately 2 × 10^6^ cells were maintained at each passage and cells were grown for approximately 14 population doublings before a final cell population was collected. Genomic DNA was purified using the Quick-DNA Miniprep Kit (Zymo Research D3024) and approximately 12 µg of genomic DNA was used as a template for sgRNA cassette amplification as described above. PCR product purification, sequencing, and screen analysis were performed as for the genome-wide CRISPR/Cas9 screens. Depletion Scores were calculated as the (NCI-237 gene log_2_ fold-change) – (TPC-1 or UTSW-354 gene log_2_ fold-change).

### GC-MS metabolomics and [U-^13^ C_6_] glucose labeling

For GC-MS metabolomics during LDH inhibitor treatment, NCI-237 cells were plated in 6 well plates at 300,000 cells/well in 3mL media and allowed to adhere overnight. The following day, glucose-free HPLM was prepared and supplemented with 5 mM [U-^13^C_6_] glucose before sterile filtration. Media was further supplemented with 2% dialyzed FBS, 1% penicillin/streptomycin, and 1X insulin-transferrin-selenium. 1 µM (R)-GNE-140, 40 nM NCGC00420737, or an equivalent volume of DMSO was added to media and mixed thoroughly. Plating media was aspirated, cells were rinsed one time with 2 mL PBS, and 2 mL media containing vehicle or LDH inhibitor was added to respective wells. Cells were incubated at 37°C and 5% CO_2_. Following incubation, cells were removed, media was rapidly aspirated, cells were rinsed one time with 4 mL ice-cold saline (0.9% sodium chloride), and 1 mL dry ice-cold extraction solvent (80:20 methanol (Thermo Fisher A456):water (Thermo Fisher W6500)) was added to cells. Cells were scraped and transferred to 1.5 mL tubes and stored at −80°C overnight. The following day, samples were centrifuged at 18,000x *g* for 10 minutes at 4°C, metabolite extracts were transferred to new tubes, and extracts were dried to completeness using a SpeedVac. Samples were derivatized by incubating with 30 µL 10 mg/mL methoxyamine hydrochloride (Sigma 226904) in anhydrous pyridine (Sigma 270970) for 15 minutes at 70°C followed by incubation with 70 µL TBDMS for 1 hour at 70°C. Derivatized samples were transferred to Agilent GC-MS autosampler vials and data acquisition was performed using an Agilent G2579A MSD coupled to an Agilent 6890 gas chromatogram. Data were analyzed using Agilent MSD Chemstation software and a MatLab script for area-under-the-curve analysis, total ion count (TIC) determination, and natural isotopomer abundance correction. For [U-^13^C_6_] glucose labeling experiments, cells were plated in 6 well plates at the following densities: NCI-237 (300,000 cells/well), XTC.UC1 (400,000 cells/well), TPC-1 (400,000 cells/well), and UTSW-354 (900,000 cells/well) in 3 mL media and allowed to adhere overnight. The following day, media containing [U-^13^C_6_] glucose was prepared as above and labeling, sample collection, sample prep, data acquisition, and data analysis were performed as above. Conditions for all GC-MS experiments were assayed in triplicate and performed one ([U-^13^C_6_] glucose labeling experiments) or two times (compound treatment for GC-MS).

### LC-MS/MS metabolomics

Snap-frozen tumors were pulverized using a dry ice- and liquid nitrogen-cooled mortar and pestle to generate a fine powder. Approximately 15-20 mg of pulverized tumor was transferred to a 2 mL tube on dry ice and 800 µL of ice-cold solvent (40:40:20 acetonitrile:methanol:water containing 0.1 M formic acid and 0.5 µM internal standard) was added to tubes. Samples were vortexed at room temperature for 10 seconds, stored on ice for 2 minutes, quenched with 70 µL 15% ammonium bicarbonate (NH_4_CO_3_), and stored on ice for 20 minutes before centrifuging at 18,000*g* for 10 minutes at 4°C. Metabolite extracts were transferred to new tubes and a second round of extraction was performed as above using 400 µL extraction solvent and 35 µL ammonium bicarbonate for quenching (Lu et al., 2017; Lu et al., 2018). Extracts from both extractions were pooled, mixed thoroughly, and transferred to LC-MS autosampler vials. LC-MS/MS data acquisition was performed as follows: chromatography was performed under HILIC conditions using a SeQuant® ZIC®-pHILIC 5 μm polymeric 150 × 2.1 mm PEEK coated HPLC column (MilliporeSigma, USA). The column temperature, sample injection volume, and flow rate were set to 45 °C, 5 μL, and 0.15 mL/min respectively. The HPLC conditions were as follows: Solvent A: 20 mM ammonium carbonate including 0.1% Ammonium hydroxide and 5 µM of Medronic acid. Solvent B: Acetonitrile. Gradient condition was 0 min: 80% B, 20 min: 20% B, 20.5 min 80% B, 34 min: 80% B. Total run time: 34 mins. Mass spectrometric analyses were performed on SCIEX QTRAP 6500+ mass spectrometer equipped with an ESI ion spray source. The ESI source was used in both positive and negative ion modes. The ion spray needle voltages used for MRM positive and negative polarity modes were set at 4800 V and −4500 V, respectively. The mass spectrometer was coupled to Shimadzu HPLC (Nexera X2 LC-30AD). The system was controlled by Analyst 1.7.2 software. Data were processed by SCIEX MultiQuant 3.0.3 software with relative quantification based on the peak area of each metabolite. TIC values were obtained using Analyst 1.7.2 software.

### NCGC00420737 measurement in plasma and tumors

Plasma was collected from potassium-EDTA-treated whole blood by centrifuging at 10,000x *g* for 10 minutes at 4°C For plasma standards and QC samples, 98 µL of blank plasma was spiked with 2 µL of standard. Drug treated samples were diluted 1:25; vehicle samples were not diluted. Standards, QC samples, and test samples of 100 µL were crashed with 200 µL of methanol containing 0.15% formic acid (final 0.1%), 3 mM ammonium acetate (final 2 mM), 75 ng/mL N-benzylbenzamide (final 50 ng/mL) internal standard (IS). The samples were vortexed for 15 seconds, incubated at room temperature for 10 minutes and centrifuged for 5 min at 16,100x *g* at 4ºC. The resulting supernatants were analyzed by LC-MS/MS. For tumors, tissues were weighed and 3 volumes of PBS were added to each tumor. Tumors were homogenized using a polytron homogenizer. 140 µL of each vehicle homogenate was combined in a single tube and diluted 1:1 with PBS to make a matrix for the standard curve. For the standards and QC samples, 98 µL of the diluted vehicle homogenate was spiked with 2 µL of standard. All samples, including vehicle were mixed 1:1 with PBS prior to analysis. Standards, QC samples, and test samples of 100 µL were crashed with 200 µL of methanol containing 0.15% formic acid (final 0.1%), 3 mM ammonium acetate (final 2 mM), 75 ng/mL N-benzylbenzamide IS (final 50 ng/mL). The samples were vortexed for 15 seconds, incubated at room temperature for 10 minutes and centrifuged for 5 min at 16,100x *g* at 4ºC. The resulting supernatants were analyzed by LC-MS/MS. Chromatography was performed using an Agilent C18 XDB column, 5 µm packing 50 × 4.6 mm size. The flow rate was set to 1.5 mL/min. The HPLC conditions were as follows: Solvent A: water + 0.1% formic acid and 2 mM ammonium acetate. Solvent B: Methanol + 0.1% formic acid and 2 mM ammonium acetate. Gradient condition was 0 – 1.5 min: 3% B; 1.5 – 2.0 min; 100% B; 2.0 – 3.5 min: 100% B; 3.5 – 3.6 min: 3% B; 3.6 – 4.5 min: 3% B. Mass spectrometric analyses were performed on SCIEX 4500 Triple Quad mass spectrometer equipped with an ESI ion spray source. The ESI source was used in positive mode with multiple-reaction monitoring.

### Histology and immunohistochemistry

NCI-237-R^PDX^ tumor fragments from vehicle- and NCGC00420737-treated animals were fixed in 10% neutral-buffered formalin and incubated at 4°C for 24 hours with gentle shaking. Fixed tumors were submitted to the UT Southwestern Histo Pathology Core for paraffin embedding and sectioning. Hematoxylin- and eosin-staining was performed by the UT Southwestern Histo Pathology Core. Ki-67 immunohistochemical staining was performed by the UT Southwestern Simmons Comprehensive Cancer Center Tissue Management Shared Resource using a BOND-RX Automated Stainer (Leica Biosystems) and human-specific anti-Ki-67 antibody (Cell Signaling Technology #9027; 1:400 dilution). Whole-slide scans were captured at 40X magnification using a Vectra Polaris slide scanner (Akoya Biosciences) and images were generated using Phenochart™ 1.1.0 software (Akoya Biosciences).

### Immunoblotting

For cell culture samples, cells were plated and allowed to adhere overnight. After treatment, media was removed, cells were washed 1 time with ice-cold PBS, and cells were lysed using Buffer A (50 mM HEPES pH 7.4, 10 mM KCl, 2 mM MgCl_2_) with 1% SDS and 1:2,000 Benzonase® nuclease (Sigma E1014). For tissue samples, pulverized tumor tissue was transferred to 1.5 mL tubes on a dry ice-cold metal block and Buffer A (50 mM HEPES pH 7.4, 10 mM KCl, 2 mM MgCl_2_) with 1% SDS and 1:2,000 Benzonase® nuclease was added to each sample. Samples were placed on a ThermoMixer at 1,000 rpm at room temperature for 15 minutes to lyse tissue. Cell and tissue debris was removed by centrifugation at 16,000x *g* for 5 minutes at room temperature and protein concentration was measured using A_280_ values determined by Cytation 5 (BioTek). Lysates were normalized in Buffer A with 1% SDS and prepared with 6X Laemmli sample buffer containing 100 mM 2-mercaptoethanol (Sigma M6250). Lysates were separated on 4-12% Bolt™ Bis-Tris gels (Invitrogen) using MES-SDS running buffer (Invitrogen B0002) and transferred to PVDF membranes using standard Towbin transfer buffer. After transfer, membranes were stained with 0.1% Ponceau S and stained membrane pictures were captured with a camera. Membranes were rinsed with TBS-0.1% Tween-20 (TBS-T) and blocked with 5% non-fat dry milk in TBS-T for 1 hour at room temperature. Membranes were rinsed with TBS-T 3 and incubated with primary antibodies overnight at 4°C. The following day, membranes were rinsed with TBS-T and incubated with secondary antibodies for 40 minutes at room temperature. Membranes were rinsed with TBS-T, incubated with Clarity Western ECL Substrate (BioRad 1705061), and exposed to X-ray film. Primary antibodies were diluted in 5% bovine serum albumin/TBS-T and used at the following concentrations: anti-phospho-AMPKα^T172^, Cell Signaling Technology #2535, 1:5,000; anti-AMPKα, Cell Signaling Technology #2532, 1:2,000; anti-FLAG M2-Peroxidase (HRP), Sigma #A8592, 1:50,000.

### Whole-exome sequencing analysis

Whole-exome sequencing data for NCI-248138-237-R [Lot# GF9] (Version 2.0.1.50.0) was downloaded from the PDMR website. Trim Galore (www.bioinformatics.babraham.ac.uk/projects/trim_galore/) was used for quality and adapter trimming. The reference genome sequences for human (hg38) and mouse (mm10) were downloaded from Illumina iGenomes (www.support.illumina.com/sequencing/sequencing_software/igenome.html). The sequencing reads were aligned to human and mouse genome sequences using Burrows-Wheeler Aligner (BWA, v0.7.17) (Li and Durbin, 2009), and contamination reads from mouse genomic DNA were removed using a custom Perl script named REMOCON (www.github.com/jiwoongbio/REMOCON). Picard (2.21.3) (www.broadinstitute.github.io/picard) was used to remove PCR duplicates and Genome Analysis Toolkit (GATK, 4.1.4.0) (DePristo et al., 2011; McKenna et al., 2010) was used to recalibrate base qualities. Calling variants and genotyping were performed using GATK HaplotypeCaller and low-quality variant calls were excluded by the following filtering thresholds: QD (Variant Confidence/Quality by Depth) < 2, FS (Phred-scaled p-value using Fisher’s exact test to detect strand bias) > 60, MQ (RMS Mapping Quality) < 40, DP (Approximate read depth) < 3, GQ (Genotype Quality) < 7. Custom Perl scripts (www.github.com/jiwoongbio/Annomen) were used to annotate variants with RefSeq (O’Leary et al., 2016) human transcripts and proteins, mitochondrial genes (NC_012920.1), dbSNP (build 151) (Sherry et al., 1999), Genome Aggregation Database (gnomAD, r3.0) (Karczewski et al., 2020), Catalogue Of Somatic Mutations In Cancer (COSMIC, v90) (www.cancer.sanger.ac.uk) (Tate et al., 2019).

### mtDNA sequencing and analysis

To lyse cells for DNA extraction, samples were digested with proteinase K (Fisher BioReagents™ BP1700-100) in digestion buffer (20 mM Tris, 100 mM NaCl, 0.5% sodium dodecyl sulfate, 10 mM EDTA, pH 7.6) at 44°C. Following digestion, samples were supplemented with additional NaCl (final concentration 2 mM) to improve DNA extraction yield. Cellular debris was pelleted via centrifugation at 14,000x *g* for 10 minutes. Total DNA was isolated from the supernatant via phenol/chloroform extraction and ethanol precipitation as previously described (Green and Sambrook, 2016; 2017). mtDNA amplification, library preparation, sequencing, and informatics analysis were performed as previously described (Wang et al., 2022b) with the following exception: trimmed reads were mapped to the *Homo sapiens* mitochondrial genome GRCh38. mtDNA coverage for each sample was as follows: NCI-237-R^PDX^ (mean: 7,239X; range: 582X – 11,608X); NCI-237^UTSW^ cell line (mean: 9,433X; range: 1018X – 14,649X); TPC-1 cell line (mean: 7,476X; range: 168X – 13,163X); UTSW-354 cell line (mean: 5,515X; range: 129X – 9,045X).

### RNA extraction and RNA-sequencing

Cells were plated in 60mm dishes in 6 mL media and allowed to adhere overnight. The following day, media was removed, cells were rinsed one time with 3 mL PBS, and 500 µL TRIzol™ (Thermo Fisher 15596026) was added to each plate. Plates were incubated at room temperature for approximately 10 minutes before several rounds of pipetting to lyse cells. The resulting lysate was transferred to a 1.5 mL tube and total RNA isolation was performed as described (Rio et al., 2010). Total RNA was submitted to BGI Genomics for poly A selection, library preparation, and sequencing. Trim Galore (www.bioinformatics.babraham.ac.uk/projects/trim_galore/) was used for quality and adapter trimming. The qualities of RNA-sequencing libraries were estimated by mapping the reads onto human transcript and ribosomal RNA sequences (Ensembl release 89) using Bowtie (v2.3.4.3) (Langmead and Salzberg, 2012). STAR (v2.7.2b) (Dobin et al., 2013) was employed to align the reads onto the human genome, SAMtools was employed to sort the alignments, and HTSeq Python package (Anders et al., 2015) was employed to count reads per gene. The values of Fragments Per Kilobase of transcript per Million mapped reads (FPKM) were calculated by a custom Perl script.

## SUPPLEMENTAL INFORMATION TITLES AND LEGENDS

**Figure S1, related to Figure 1.**
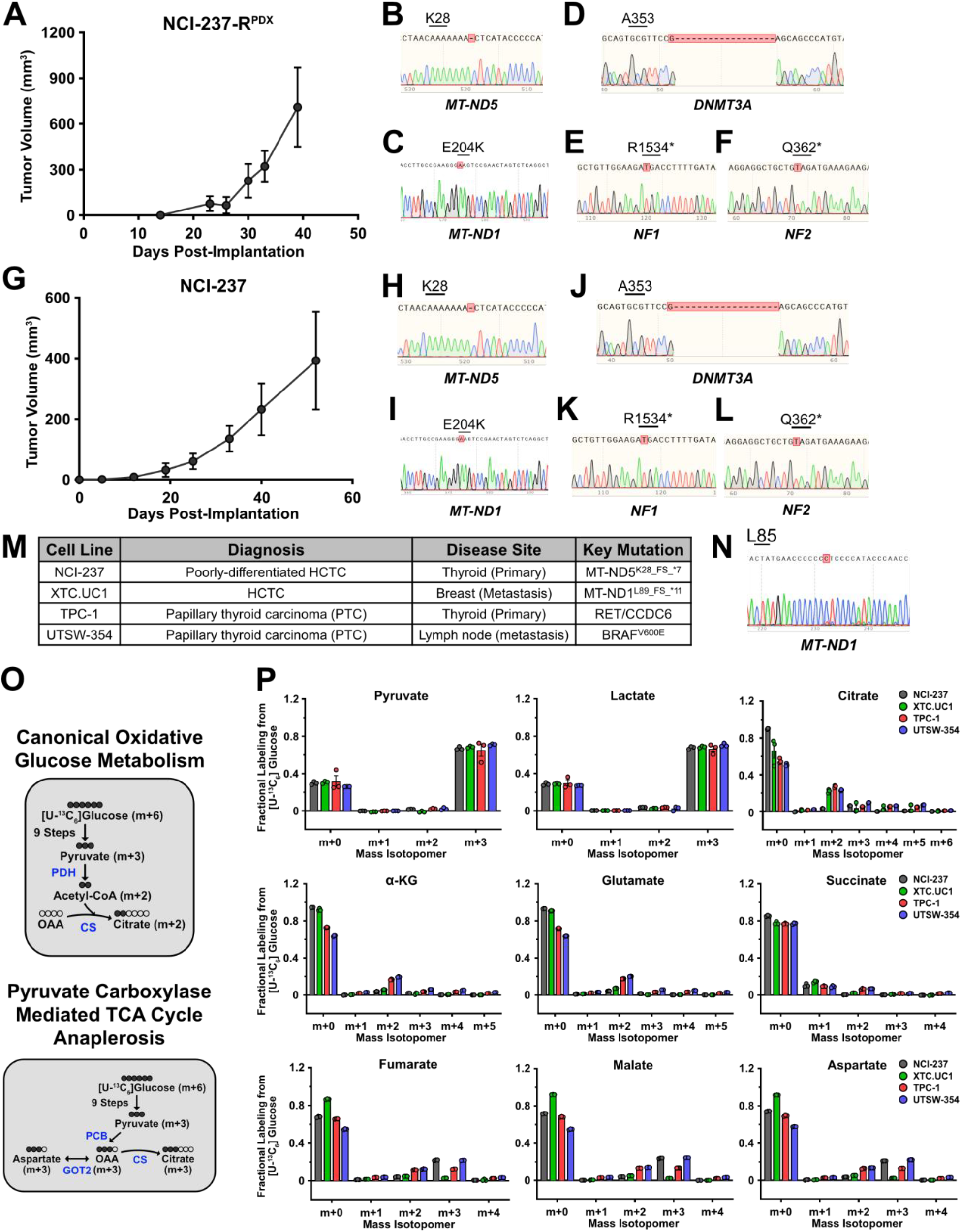
A) Tumor volume of NCI-237-R^PDX^ implanted subcutaneously in immunodeficient mice. Data are plotted as mean ± SD of 4 tumors. B) Sanger sequencing trace of NCI-237-R^PDX^ PCR-amplified *MT-ND5*. C) Sanger sequencing trace of NCI-237-R^PDX^ PCR-amplified *MT-ND1*. D) Sanger sequencing trace of NCI-237-R^PDX^ PCR-amplified *DNMT3A*. E) Sanger sequencing trace of NCI-237-R^PDX^ PCR-amplified *NF1*. F) Sanger sequencing trace of NCI-237-R^PDX^ PCR-amplified *NF2*. G) Tumor volume of NCI-237 cells injected subcutaneously in immunodeficient mice. Data are plotted as mean ± SD of 4 tumors. H) Sanger sequencing trace of NCI-237^UTSW^ PCR-amplified *MT-ND5*. I) Sanger sequencing trace of NCI-237^UTSW^ PCR-amplified *MT-ND1*. J) Sanger sequencing trace of NCI-237^UTSW^ PCR-amplified *DNMT3A*. K) Sanger sequencing trace of NCI-237^UTSW^ PCR-amplified *NF1*. L) Sanger sequencing trace of NCI-237^UTSW^ PCR-amplified *NF2*. M) Table outlining cell lines used in this study. N) Sanger sequencing trace of XTC.UC1 PCR-amplified *MT-ND1*. O) Schematic for labeling of key central carbon metabolites from [U-^13^C_6_] glucose under canonical oxidative conditions (*top*) or pyruvate carboxylase-mediated TCA cycle anaplerosis (*bottom*). PDH = pyruvate dehydrogenase; CS = citrate synthase; PCB = pyruvate carboxylase; GOT2 = glutamic-oxaloacetic transaminase 2; OAA = oxaloacetate. P) Mass isotopomer abundance for the indicated metabolites in cells cultured with [U-^13^C_6_] glucose for 6 hours. Data are plotted as mean ± SEM of 3 replicates. α-KG = alpha-ketoglutarate.

**Figure S2, related to Figure 2.**
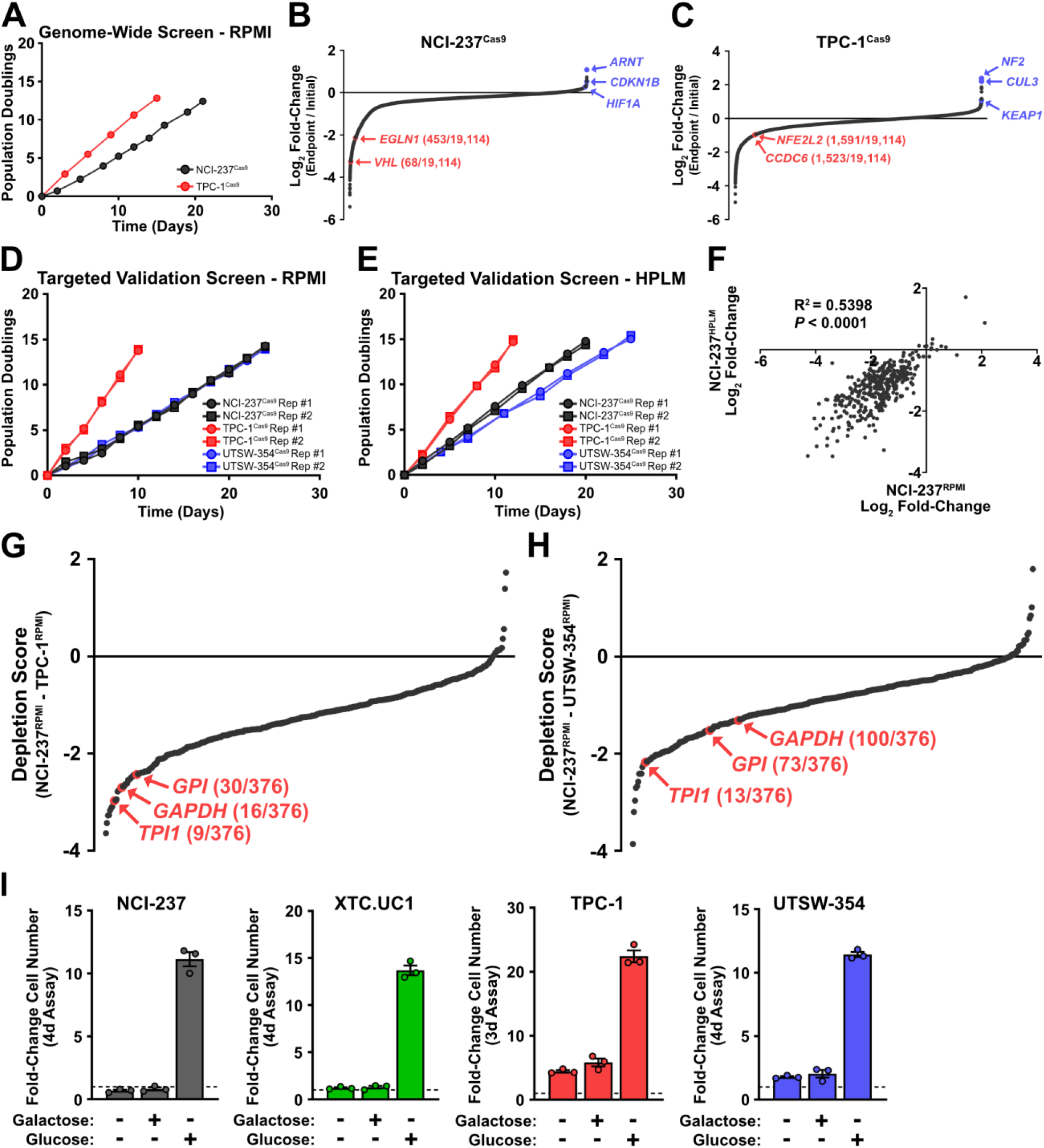
A) Growth curves of NCI-237^RPMI^ and TPC-1^RPMI^ in primary genome-wide CRISPR/Cas9 knockout screen. B) Plot of genes for NCI-237^RPMI^ based on median log_2_ fold-change in sgRNA abundance. C) Plot of genes for TPC-1^RPMI^ based on median log_2_ fold-change in sgRNA abundance. Enriched genes are highlighted in blue; depleted genes are highlighted in red. D) Growth curves of NCI-237^RPMI^, TPC-1^RPMI^, and UTSW-354^RPMI^ replicates (*n* = 2) in targeted validation screen. E) Growth curves of NCI-237^HPLM^, TPC-1^HPLM^, and UTSW-354^HPLM^ replicates (*n* = 2) in targeted validation screen. F) Plot of gene log_2_ fold-change values for NCI-237^RPMI^ and NCI-237^HPLM^ validation screens. Data reflect median log_2_ fold-change in sgRNA abundance based on 2 replicates for each condition. G) Plot of genes from validation screen based on Depletion Score comparing NCI-237^RPMI^ and TPC-1^RPMI^. H) Plot of genes from validation screen based on Depletion Score comparing NCI-237^RPMI^ and UTSW-354^RPMI^. Depletion Score for G) and H) are calculated as [(NCI-237^RPMI^ gene log_2_ fold-change) – (TPC-1^RPMI^/UTSW-354^RPMI^ gene log_2_ fold-change)]. Gene log_2_ fold-change values reflect median log_2_ fold-change in sgRNA abundance based on 2 replicates for each cell line. I) Fold-change in cell number of indicated cell lines in media lacking glucose or galactose, media containing 5 mM galactose, or media containing 5 mM glucose. Data are plotted as mean ± SEM of 3 replicates. Dashed line = 1.

**Figure S3, related to Figure 3.**
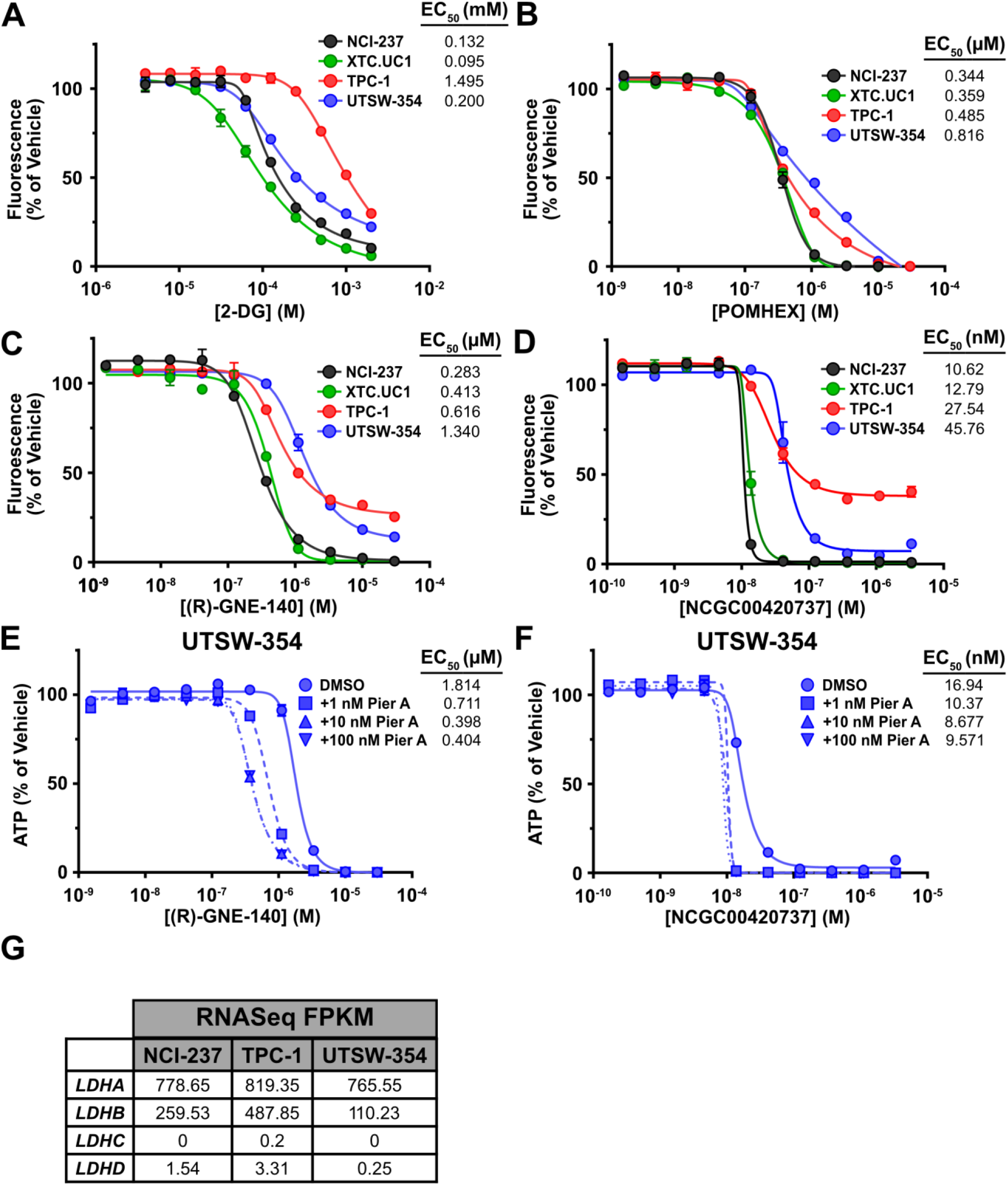
A) Viability assay for cells treated with 2-DG for 3 days. B) Viability assay for cells treated with POMHEX for 3 days. C) Viability assay for cells treated with (R)-GNE-140 for 3 days. D) Viability assay for cells treated with NCGC00420737 for 3 days. E) Viability assay for UTSW-354 cotreated with (R)-GNE-140 and indicated doses of piericidin A for 3 days. F) Viability assay for UTSW-354 cotreated with NCGC00420737 and indicated doses of piericidin A for 3 days. Data are plotted as mean ± SEM of 2 replicates. G) Table of RNA-Seq FPKM values for *LDH* genes in indicated cell lines.

**Figure S4, related to Figure 4.**
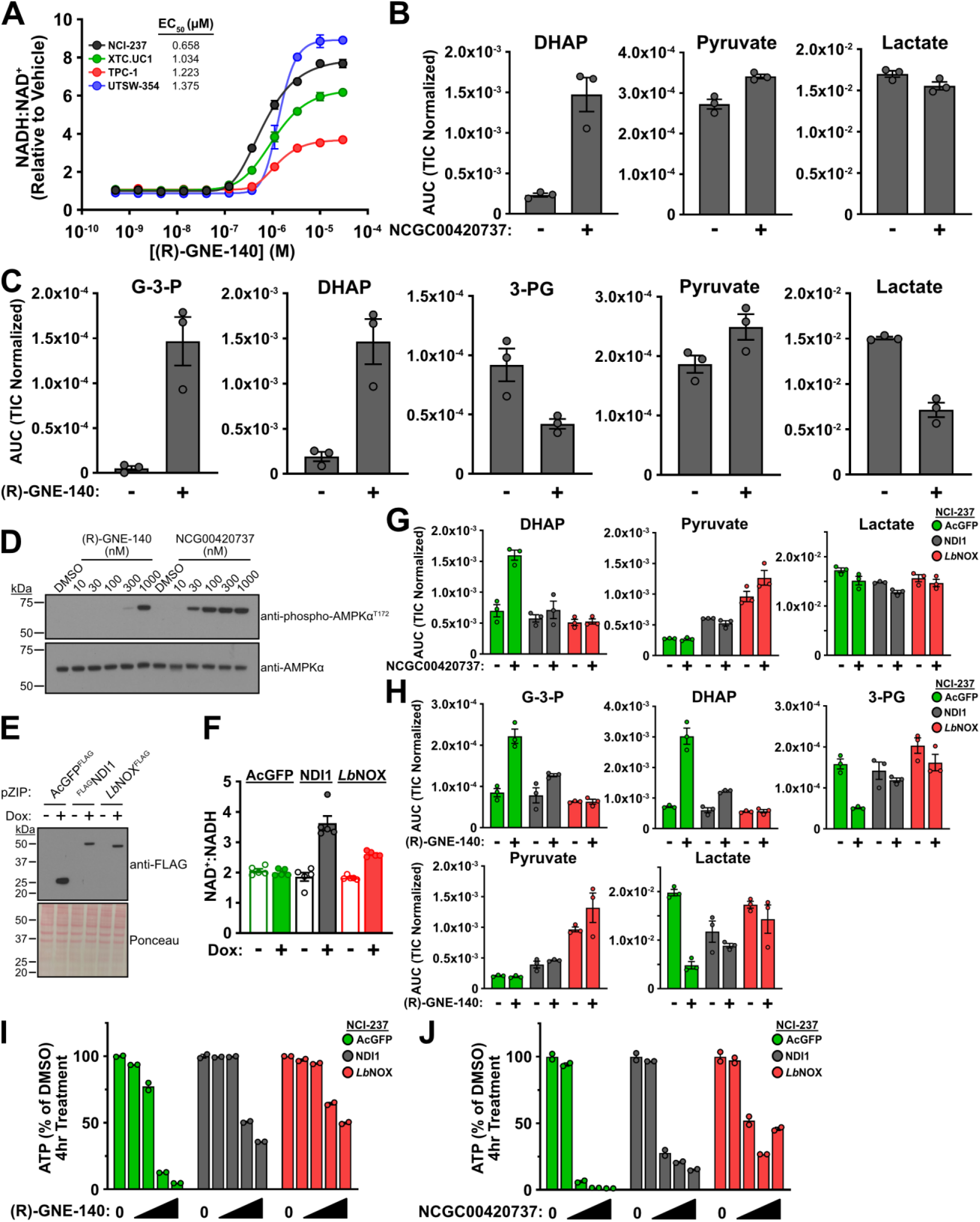
A) NADH:NAD^+^ ratio in cells treated with (R)-GNE-140 for 4 hours; *n* = 2 replicates. B) Metabolite levels in NCI-237 cells treated with 40 nM NCGC00420737 for 4 hours; *n* = 3 replicates. DHAP = dihydroxyacetone phosphate. C) Metabolite levels in NCI-237 cells treated with 1 μM (R)-GNE-140 for 4 hours; *n* = 3 replicates. G-3-P = glyceraldehyde-3-phosphate; DHAP = dihydroxyacetone phosphate; 3-PG = 3-phosphoglycerate. D) Immunoblot for phospho-AMPKα^T172^ and total AMPKα in NCI-237 cells treated with indicated doses of (R)-GNE-140 and NCGC00420737 for 4 hours. E) Immunoblot for FLAG epitope tag in NCI-237 cells with inducible expression of indicated constructs. Cells were exposed to 100 ng/mL doxycycline for 24 hours prior to collection. F) NAD^+^:NADH ratio in NCI-237 cells with inducible expression of indicated constructs. Cells were exposed to 100 ng/mL doxycycline for 24 hours prior to collection; *n* = 5 replicates. G) Metabolite levels in NCI-237 cells expressing AcGFP, NDI1, or *Lb*NOX treated with 40 nM NCGC00420737 for 4 hours; *n* = 3 replicates. DHAP = dihydroxyacetone phosphate. H) Metabolite levels in NCI-237 cells expressing AcGFP, NDI1, or *Lb*NOX treated with 1 μM (R)-GNE-140 for 4 hours; *n* = 3 replicates. G-3-P = glyceraldehyde-3-phosphate; DHAP = dihydroxyacetone phosphate; 3-PG = 3-phosphoglycerate. I) ATP levels in cells treated with (R)-GNE-140 for 4 hours; *n* = 2 replicates. Concentrations are 0.041, 0.37, 3.33, and 30 µM (R)-GNE-140. J) ATP levels in cells treated with NCGC00420737 for 4 hours; *n* = 2 replicates. Concentrations are 0.014, 0.123, 1.11, and 10 µM NCGC00420737. Data are plotted as mean ± SEM of indicated number of replicates.

**Figure S5, related to Figure 5.**
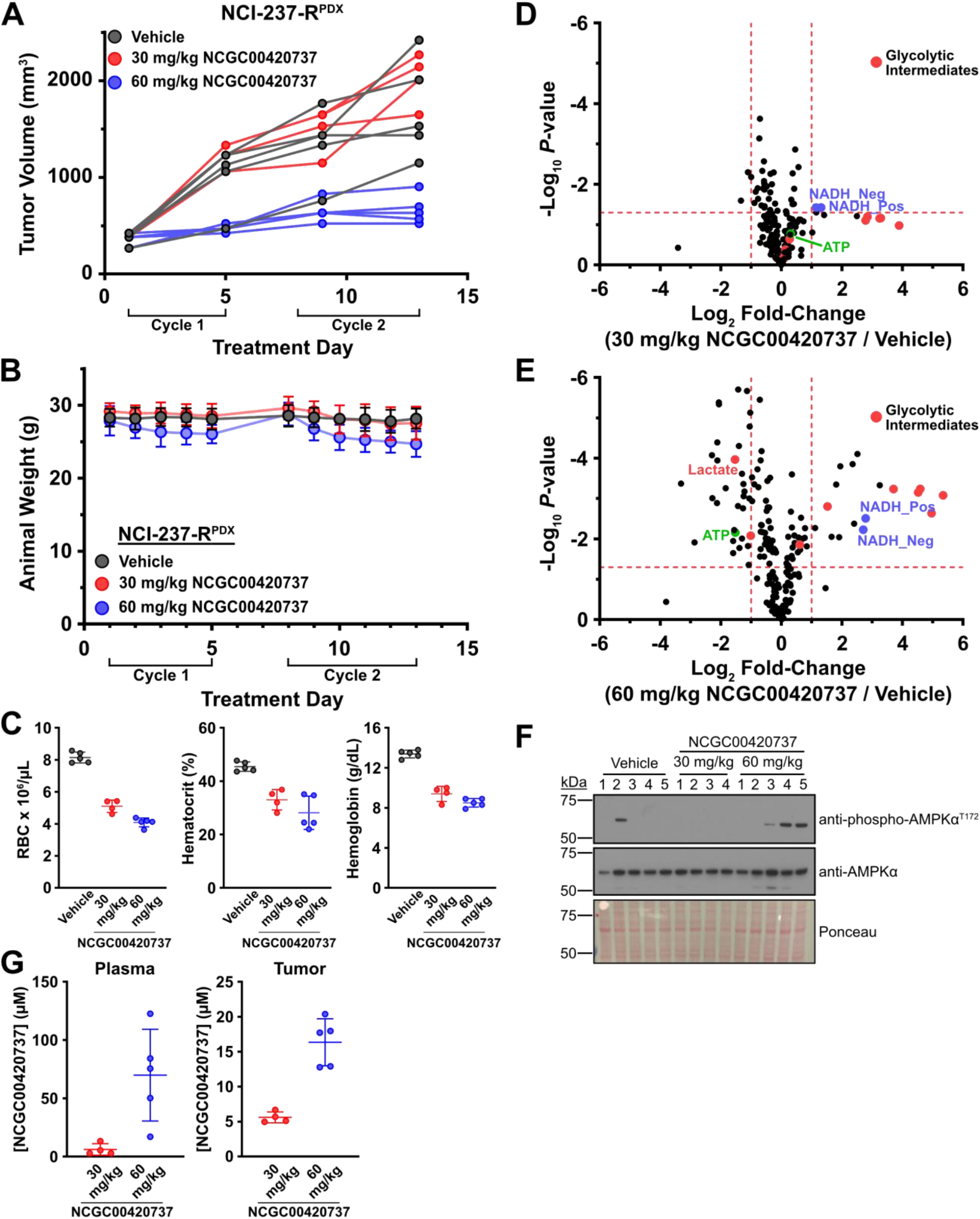
A) Individual tumor volumes for NCI-237-R^PDX^ xenografts treated with vehicle (*n* = 5), 30 mg/kg NCGC00420737 (*n* = 4), or 60 mg/kg NCGC00420737 (*n* = 5). Vehicle or compound was administered once daily via jugular vein catheter for two treatment cycles (5 days on, 2 days off). A final compound administration was performed on Day 6 of Cycle 2 and animals were sacrificed 1 hour after receiving compound. B) Animal weight for NCI-237-R^PDX^-bearing immunodeficient mice treated with vehicle (*n* = 5), 30 mg/kg NCGC00420737 (*n* = 4), or 60 mg/kg NCGC00420737 (n = 5). C) Red blood cell (left), hematocrit (center), and hemoglobin (right) levels in EDTA-treated whole blood. Blood was collected 1 hour after final administration of compound. D) Volcano plot comparing metabolite levels in NCI-237-R^PDX^ tumors treated with vehicle and 30 mg/kg NCGC00420737. Metabolite levels were determined using LC-MS/MS. Horizontal dashed line = 1.3 (*P*-value ≅ 0.05); vertical dashed lines = −1 and 1. E) Volcano plot comparing metabolite levels in NCI-237-R^PDX^ tumors treated with vehicle and 60 mg/kg NCGC00420737. Metabolite levels were determined using LC-MS/MS. Horizontal dashed line = 1.3 (*P*-value ≅ 0.05); vertical dashed lines = −1 and 1. *P*-values in D) and E) were determined using two-sided *T*-test assuming unequal variance. F) Immunoblot for phospho-AMPKα^T172^ and total AMPKα in NCI-237-R^PDX^ tumors. G) Plasma and tumor concentrations of NCGC00420737 determined by LC-MS/MS. Blood and tumors were collected 1 hour after final administration of compound. Data are plotted as mean ± SD of indicated number of replicates.

## Notes

### Competing Interest Statement

The authors have declared no competing interest.

## REFERENCES

Anders, S., Pyl, P.T., and Huber, W. (2015). HTSeq--a Python framework to work with high-throughput sequencing data. Bioinformatics 31, 166–169. 10.1093/bioinformatics/btu638.

Arroyo, J.D., Jourdain, A.A., Calvo, S.E., Ballarano, C.A., Doench, J.G., Root, D.E., and Mootha, V.K. (2016). A Genome-wide CRISPR Death Screen Identifies Genes Essential for Oxidative Phosphorylation. Cell Metab 24, 875–885. 10.1016/j.cmet.2016.08.017.

Birsoy, K., Possemato, R., Lorbeer, F.K., Bayraktar, E.C., Thiru, P., Yucel, B., Wang, T., Chen, W.W., Clish, C.B., and Sabatini, D.M. (2014). Metabolic determinants of cancer cell sensitivity to glucose limitation and biguanides. Nature 508, 108–112. 10.1038/nature13110.

Birsoy, K., Wang, T., Chen, W.W., Freinkman, E., Abu-Remaileh, M., and Sabatini, D.M. (2015). An Essential Role of the Mitochondrial Electron Transport Chain in Cell Proliferation Is to Enable Aspartate Synthesis. Cell 162, 540–551. 10.1016/j.cell.2015.07.016.

Bonora, E., Porcelli, A.M., Gasparre, G., Biondi, A., Ghelli, A., Carelli, V., Baracca, A., Tallini, G., Martinuzzi, A., Lenaz, G., et al. (2006). Defective oxidative phosphorylation in thyroid oncocytic carcinoma is associated with pathogenic mitochondrial DNA mutations affecting complexes I and III. Cancer Res 66, 6087–6096. 10.1158/0008-5472.CAN-06-0171.

Boudreau, A., Purkey, H.E., Hitz, A., Robarge, K., Peterson, D., Labadie, S., Kwong, M., Hong, R., Gao, M., Del Nagro, C., et al. (2016). Metabolic plasticity underpins innate and acquired resistance to LDHA inhibition. Nat Chem Biol 12, 779–786. 10.1038/nchembio.2143.

Cantor, J.R., Abu-Remaileh, M., Kanarek, N., Freinkman, E., Gao, X., Louissaint, A., Jr., Lewis, C.A., and Sabatini, D.M. (2017). Physiologic Medium Rewires Cellular Metabolism and Reveals Uric Acid as an Endogenous Inhibitor of UMP Synthase. Cell 169, 258–272 e217. 10.1016/j.cell.2017.03.023.

Corver, W.E., Demmers, J., Oosting, J., Sahraeian, S., Boot, A., Ruano, D., Wezel, T.V., and Morreau, H. (2018). ROS-induced near-homozygous genomes in thyroid cancer. Endocr Relat Cancer 25, 83–97. 10.1530/ERC-17-0288.

Courtney, K.D., Bezwada, D., Mashimo, T., Pichumani, K., Vemireddy, V., Funk, A.M., Wimberly, J., McNeil, S.S., Kapur, P., Lotan, Y., et al. (2018). Isotope Tracing of Human Clear Cell Renal Cell Carcinomas Demonstrates Suppressed Glucose Oxidation In Vivo. Cell Metab 28, 793–800 e792. 10.1016/j.cmet.2018.07.020.

Crooks, D.R., Maio, N., Lang, M., Ricketts, C.J., Vocke, C.D., Gurram, S., Turan, S., Kim, Y.Y., Cawthon, G.M., Sohelian, F., et al. (2021). Mitochondrial DNA alterations underlie an irreversible shift to aerobic glycolysis in fumarate hydratase-deficient renal cancer. Sci Signal 14. 10.1126/scisignal.abc4436.

DePristo, M.A., Banks, E., Poplin, R., Garimella, K.V., Maguire, J.R., Hartl, C., Philippakis, A.A., del Angel, G., Rivas, M.A., Hanna, M., et al. (2011). A framework for variation discovery and genotyping using next-generation DNA sequencing data. Nat Genet 43, 491–498. 10.1038/ng.806.

Ditta, G., Soderberg, K., Landy, F., and Scheffler, I.E. (1976). The selection of Chinese hamster cells deficient in oxidative energy metabolism. Somatic Cell Genet 2, 331–344. 10.1007/BF01538838.

Dobin, A., Davis, C.A., Schlesinger, F., Drenkow, J., Zaleski, C., Jha, S., Batut, P., Chaisson, M., and Gingeras, T.R. (2013). STAR: ultrafast universal RNA-seq aligner. Bioinformatics 29, 15–21. 10.1093/bioinformatics/bts635.

Ganly, I., Liu, E.M., Kuo, F., Makarov, V., Dong, Y., Park, J., Gong, Y., Gorelick, A.N., Knauf, J.A., Benedetti, E., et al. (2022). Mitonuclear genotype remodels the metabolic and microenvironmental landscape of Hurthle cell carcinoma. Sci Adv 8, eabn9699. 10.1126/sciadv.abn9699.

Ganly, I., Makarov, V., Deraje, S., Dong, Y., Reznik, E., Seshan, V., Nanjangud, G., Eng, S., Bose, P., Kuo, F., et al. (2018). Integrated Genomic Analysis of Hurthle Cell Cancer Reveals Oncogenic Drivers, Recurrent Mitochondrial Mutations, and Unique Chromosomal Landscapes. Cancer Cell 34, 256–270 e255. 10.1016/j.ccell.2018.07.002.

Ganly, I., and McFadden, D.G. (2019). Short Review: Genomic Alterations in Hurthle Cell Carcinoma. Thyroid 29, 471–479. 10.1089/thy.2019.0088.

Gopal, R.K., Kubler, K., Calvo, S.E., Polak, P., Livitz, D., Rosebrock, D., Sadow, P.M., Campbell, B., Donovan, S.E., Amin, S., et al. (2018). Widespread Chromosomal Losses and Mitochondrial DNA Alterations as Genetic Drivers in Hurthle Cell Carcinoma. Cancer Cell 34, 242–255 e245. 10.1016/j.ccell.2018.06.013.

Green, M.R., and Sambrook, J. (2016). Precipitation of DNA with ethanol. Cold Spring Harbor Protocols 2016, pdb. prot093377.

Green, M.R., and Sambrook, J. (2017). Isolation of high-molecular-weight DNA using organic solvents. Cold Spring Harbor Protocols 2017, pdb. prot093450.

Hawley, S.A., Davison, M., Woods, A., Davies, S.P., Beri, R.K., Carling, D., and Hardie, D.G. (1996). Characterization of the AMP-activated protein kinase kinase from rat liver and identification of threonine 172 as the major site at which it phosphorylates AMP-activated protein kinase. J Biol Chem 271, 27879–27887. 10.1074/jbc.271.44.27879.

Joung, J., Konermann, S., Gootenberg, J.S., Abudayyeh, O.O., Platt, R.J., Brigham, M.D., Sanjana, N.E., and Zhang, F. (2017). Genome-scale CRISPR-Cas9 knockout and transcriptional activation screening. Nat Protoc 12, 828–863. 10.1038/nprot.2017.016.

Karczewski, K.J., Francioli, L.C., Tiao, G., Cummings, B.B., Alfoldi, J., Wang, Q., Collins, R.L., Laricchia, K.M., Ganna, A., Birnbaum, D.P., et al. (2020). The mutational constraint spectrum quantified from variation in 141,456 humans. Nature 581, 434–443. 10.1038/s41586-020-2308-7.

King, M.P., and Attardi, G. (1989). Human cells lacking mtDNA: repopulation with exogenous mitochondria by complementation. Science 246, 500–503. 10.1126/science.2814477.

Koppenol, W.H., Bounds, P.L., and Dang, C.V. (2011). Otto Warburg’s contributions to current concepts of cancer metabolism. Nat Rev Cancer 11, 325–337. 10.1038/nrc3038.

Langmead, B., and Salzberg, S.L. (2012). Fast gapped-read alignment with Bowtie 2. Nat Methods 9, 357–359. 10.1038/nmeth.1923.

Li, H., and Durbin, R. (2009). Fast and accurate short read alignment with Burrows-Wheeler transform. Bioinformatics 25, 1754–1760. 10.1093/bioinformatics/btp324.

Lin, Y.H., Satani, N., Hammoudi, N., Yan, V.C., Barekatain, Y., Khadka, S., Ackroyd, J.J., Georgiou, D.K., Pham, C.D., Arthur, K., et al. (2020). An enolase inhibitor for the targeted treatment of ENO1-deleted cancers. Nat Metab 2, 1413–1426. 10.1038/s42255-020-00313-3.

Lu, W., Su, X., Klein, M.S., Lewis, I.A., Fiehn, O., and Rabinowitz, J.D. (2017). Metabolite Measurement: Pitfalls to Avoid and Practices to Follow. Annu Rev Biochem 86, 277–304. 10.1146/annurev-biochem-061516-044952.

Lu, W., Wang, L., Chen, L., Hui, S., and Rabinowitz, J.D. (2018). Extraction and Quantitation of Nicotinamide Adenine Dinucleotide Redox Cofactors. Antioxid Redox Signal 28, 167–179. 10.1089/ars.2017.7014.

Luengo, A., Gui, D.Y., and Vander Heiden, M.G. (2017). Targeting Metabolism for Cancer Therapy. Cell Chem Biol 24, 1161–1180. 10.1016/j.chembiol.2017.08.028.

McFadden, D.G., and Sadow, P.M. (2021). Genetics, Diagnosis, and Management of Hurthle Cell Thyroid Neoplasms. Front Endocrinol (Lausanne) 12, 696386. 10.3389/fendo.2021.696386.

McKenna, A., Hanna, M., Banks, E., Sivachenko, A., Cibulskis, K., Kernytsky, A., Garimella, K., Altshuler, D., Gabriel, S., Daly, M., and DePristo, M.A. (2010). The Genome Analysis Toolkit: a MapReduce framework for analyzing next-generation DNA sequencing data. Genome Res 20, 1297–1303. 10.1101/gr.107524.110.

Mullen, A.R., Wheaton, W.W., Jin, E.S., Chen, P.H., Sullivan, L.B., Cheng, T., Yang, Y., Linehan, W.M., Chandel, N.S., and DeBerardinis, R.J. (2011). Reductive carboxylation supports growth in tumour cells with defective mitochondria. Nature 481, 385–388. 10.1038/nature10642.

O’Leary, N.A., Wright, M.W., Brister, J.R., Ciufo, S., Haddad, D., McVeigh, R., Rajput, B., Robbertse, B., Smith-White, B., Ako-Adjei, D., et al. (2016). Reference sequence (RefSeq) database at NCBI: current status, taxonomic expansion, and functional annotation. Nucleic Acids Res 44, D733–745. 10.1093/nar/gkv1189.

Oshima, N., Ishida, R., Kishimoto, S., Beebe, K., Brender, J.R., Yamamoto, K., Urban, D., Rai, G., Johnson, M.S., Benavides, G., et al. (2020). Dynamic Imaging of LDH Inhibition in Tumors Reveals Rapid In Vivo Metabolic Rewiring and Vulnerability to Combination Therapy. Cell Rep 30, 1798–1810 e1794. 10.1016/j.celrep.2020.01.039.

Patgiri, A., Skinner, O.S., Miyazaki, Y., Schleifer, G., Marutani, E., Shah, H., Sharma, R., Goodman, R.P., To, T.L., Robert Bao, X., et al. (2020). An engineered enzyme that targets circulating lactate to alleviate intracellular NADH:NAD(+) imbalance. Nat Biotechnol 38, 309–313. 10.1038/s41587-019-0377-7.

Rai, G., Brimacombe, K.R., Mott, B.T., Urban, D.J., Hu, X., Yang, S.M., Lee, T.D., Cheff, D.M., Kouznetsova, J., Benavides, G.A., et al. (2017). Discovery and Optimization of Potent, Cell-Active Pyrazole-Based Inhibitors of Lactate Dehydrogenase (LDH). J Med Chem 60, 9184–9204. 10.1021/acs.jmedchem.7b00941.

Rai, G., Urban, D.J., Mott, B.T., Hu, X., Yang, S.M., Benavides, G.A., Johnson, M.S., Squadrito, G.L., Brimacombe, K.R., Lee, T.D., et al. (2020). Pyrazole-Based Lactate Dehydrogenase Inhibitors with Optimized Cell Activity and Pharmacokinetic Properties. J Med Chem 63, 10984–11011. 10.1021/acs.jmedchem.0c00916.

Rio, D.C., Ares, M., Jr., Hannon, G.J., and Nilsen, T.W. (2010). Purification of RNA using TRIzol (TRI reagent). Cold Spring Harb Protoc 2010, pdb prot5439. 10.1101/pdb.prot5439.

Robinson, B.H., Petrova-Benedict, R., Buncic, J.R., and Wallace, D.C. (1992). Nonviability of cells with oxidative defects in galactose medium: a screening test for affected patient fibroblasts. Biochem Med Metab Biol 48, 122–126. 10.1016/0885-4505(92)90056-5.

Rossiter, N.J., Huggler, K.S., Adelmann, C.H., Keys, H.R., Soens, R.W., Sabatini, D.M., and Cantor, J.R. (2021). CRISPR screens in physiologic medium reveal conditionally essential genes in human cells. Cell Metab 33, 1248–1263 e1249. 10.1016/j.cmet.2021.02.005.

Salabei, J.K., Gibb, A.A., and Hill, B.G. (2014). Comprehensive measurement of respiratory activity in permeabilized cells using extracellular flux analysis. Nat Protoc 9, 421–438. 10.1038/nprot.2014.018.

Seo, B.B., Kitajima-Ihara, T., Chan, E.K., Scheffler, I.E., Matsuno-Yagi, A., and Yagi, T. (1998). Molecular remedy of complex I defects: rotenone-insensitive internal NADH-quinone oxidoreductase of Saccharomyces cerevisiae mitochondria restores the NADH oxidase activity of complex I-deficient mammalian cells. Proc Natl Acad Sci U S A 95, 9167–9171. 10.1073/pnas.95.16.9167.

Sherry, S.T., Ward, M., and Sirotkin, K. (1999). dbSNP-database for single nucleotide polymorphisms and other classes of minor genetic variation. Genome Res 9, 677–679.

Stein, S.C., Woods, A., Jones, N.A., Davison, M.D., and Carling, D. (2000). The regulation of AMP-activated protein kinase by phosphorylation. Biochem J 345 Pt 3, 437–443.

Sullivan, L.B., Gui, D.Y., Hosios, A.M., Bush, L.N., Freinkman, E., and Vander Heiden, M.G. (2015). Supporting Aspartate Biosynthesis Is an Essential Function of Respiration in Proliferating Cells. Cell 162, 552–563. 10.1016/j.cell.2015.07.017.

Tate, J.G., Bamford, S., Jubb, H.C., Sondka, Z., Beare, D.M., Bindal, N., Boutselakis, H., Cole, C.G., Creatore, C., Dawson, E., et al. (2019). COSMIC: the Catalogue Of Somatic Mutations In Cancer. Nucleic Acids Res 47, D941–D947. 10.1093/nar/gky1015.

Titov, D.V., Cracan, V., Goodman, R.P., Peng, J., Grabarek, Z., and Mootha, V.K. (2016). Complementation of mitochondrial electron transport chain by manipulation of the NAD+/NADH ratio. Science 352, 231–235. 10.1126/science.aad4017.

Trefts, E., and Shaw, R.J. (2021). AMPK: restoring metabolic homeostasis over space and time. Mol Cell 81, 3677–3690. 10.1016/j.molcel.2021.08.015.

Wang, X., Shelton, S.D., Bordieanu, B., Frank, A.R., Yi, Y., Venigalla, S.S.K., Gu, Z., Lenser, N.P., Glogauer, M., Chandel, N.S., et al. (2022a). Scinderin promotes fusion of electron transport chain dysfunctional muscle stem cells with myofibers. Nat Aging 2, 155–169. 10.1038/s43587-021-00164-x.

Wang, X., Shelton, S.D., Bordieanu, B., Frank, A.R., Yi, Y., Venigalla, S.S.K., Gu, Z., Lesner, N.P., Glogauer, M., and Chandel, N.S. (2022b). Scinderin promotes fusion of electron transport chain dysfunctional muscle stem cells with myofibers. Nature aging 2, 155–169.

Warburg, O. (1956). On the origin of cancer cells. Science 123, 309–314. 10.1126/science.123.3191.309.

Wick, A.N., Drury, D.R., Nakada, H.I., and Wolfe, J.B. (1957). Localization of the primary metabolic block produced by 2-deoxyglucose. J Biol Chem 224, 963–969.

Yeung, C., Gibson, A.E., Issaq, S.H., Oshima, N., Baumgart, J.T., Edessa, L.D., Rai, G., Urban, D.J., Johnson, M.S., Benavides, G.A., et al. (2019). Targeting Glycolysis through Inhibition of Lactate Dehydrogenase Impairs Tumor Growth in Preclinical Models of Ewing Sarcoma. Cancer Res 79, 5060–5073. 10.1158/0008-5472.CAN-19-0217.

Yuan, Y., Ju, Y.S., Kim, Y., Li, J., Wang, Y., Yoon, C.J., Yang, Y., Martincorena, I., Creighton, C.J., Weinstein, J.N., et al. (2020). Comprehensive molecular characterization of mitochondrial genomes in human cancers. Nat Genet 52, 342–352. 10.1038/s41588-019-0557-x.

Zielke, A., Tezelman, S., Jossart, G.H., Wong, M., Siperstein, A.E., Duh, Q.Y., and Clark, O.H. (1998). Establishment of a highly differentiated thyroid cancer cell line of Hurthle cell origin. Thyroid 8, 475–483. 10.1089/thy.1998.8.475.

